# Long term exposure of human gut microbiota with high and low emulsifier sensitivity to soy lecithin in M-SHIME model

**DOI:** 10.1101/2021.12.16.472798

**Authors:** Lisa Miclotte, Ellen De Paepe, Qiqiong Li, John Van Camp, Andreja Rajkovic, Tom Van de Wiele

## Abstract

In the context of the potential health hazards related to food processing, dietary emulsifiers have been shown to alter the structure and function of the gut microbial community, both *in vivo* and *in vitro*. In mouse models, these emulsifier exposed gut microbiota were shown to contribute to gut inflammation. Several knowledge gaps remain to be addressed though. As such, the impact from a longer timeframe of exposure on the gut microbiota is not known and interindividual variability in microbiome response needs to be measured.

To answer these research questions, in this study the faecal microbiota from two individuals, previously selected for high and low emulsifier sensitivity, were exposed to two concentrations of soy lecithin during a 7 day treatment phase in the dynamic mucosal simulator of the human intestinal microbial ecosystem (M-SHIME). The results showed mild effects from soy lecithin on the composition and functionality of these microbial communities, which depended on the original microbial composition. The effects also mostly levelled off after 3 days of exposure. The emulsifier sensitivity for which the microbiota were selected, was preserved. Some potentially concerning effects were also registered: butyrate levels, positively correlating with *Faecalibacterium* abundance, were lowered by soy lecithin. Also the abundance of the beneficial *Bifidobacterium* genus was lowered, while the abundance of the notorious unclassified Enterobacteriaceae was increased. Within the family of the unclassified Lachnospiraceae, several genera were either suppressed or stimulated.

The effects that these microbial alterations would have on a living host is not yet certain, especially given the fact that large fractions of soy lecithin’s constituents can be absorbed. Nevertheless, choline and phosphatidylcholine, both primary and absorbable constituents of soy lecithin, have recently been linked to cardiovascular disease via the generation of TMA by the gut microbiota. Further studies that validate our findings and link them to potential health outcomes are thus justified.

## 1. Introduction

Metabolic syndrome is defined as a set of conditions/symptoms that are risk factors for the development of the top most lethal diseases worldwide, notably cardiovascular disease (CVD) and diabetes [1]. In general, overweight or obesity and insulin resistance have been marked as the most important risk factors [1]. Cardiovascular events (stroke and coronary artery disease) cause 32% of the world’s annual deaths [2] and lifestyle factors such as diet, exercise, smoking, alcohol abuse and pollution play an important role in their development [2]. Regarding diet, the role of salt, saturated and trans-fats and sugar with CVD has been researched extensively. However, none of these factors could explain the full picture, nor has the emergence of food products reduced in these components solved the issues. Consequently, a more overarching view of the relationship between nutrition and health has been proposed, in which the overconsumption of highly processed food products is being scrutinized [3]. The use of food additives is a hallmark of highly processed food products [4]. Emulsifiers and stabilizers are one class of these food additives that have recently been investigated for their purported destabilizing effects on the gut microbiota [5–7]. As such, chassaing et al have found in a number of studies that TWEEN80 and CMC can contribute to gut inflammation by causing microbial translocation across the intestinal mucus layer [8–10].

The human gut microbiota consists of a vast set of microorganisms that resides in the human gastrointestinal tract [11,12]. It has been evidenced that it plays a role in or is influenced by obesity and metabolic aberrations in the host [13–16]. It is also known that nutrition has a profound and rapid effect on its composition and functionality [17,18].

The emulsifier soy lecithin is an extract of soybeans that is widely used in food and beverage industry, animal feed, pharmaceuticals, cosmetics and other industrial applications [19]. In food products it is used for instance in bakery products, confectionary, ice cream, creamers and margarines [20]. Its function is to modify viscosity, capture flavours and moisture, soften doughs, etc [21]. Soy lecithin generally consists of about 60% phospholipids (phosphatidylcholine, -ethylamine, -serine and -inositol), about 35% triglycerides and a small fraction of sterols and carbohydrates [20].

We previously investigated the impact of soy lecithin on the gut microbiota from 10 human donors *in vitro*, and found that soy lecithin has the potential to stimulate propionate and acetate production, while decreasing butyrate production. The extent of these effects depended on the individual’s microbiome. In those experiments, the exposure to the emulsifier was quite short (48h).

In real life situations, individuals may be exposed to soy lecithin repeatedly over time. Perhaps even on a daily basis. It is unknown how the gut microbiota reacts to such chronic emulsifier exposure in terms of composition and functionality. Here, we therefore prolonged the exposure time up to 7 days and utilized the simulator of the human intestinal microbial ecosystem (SHIME), a validated dynamic *in vitro* model of the human gut. From our preliminary screening of 10 human individuals, we selected the two donors whose microbiota responded the most and the least to the supplemented emulsifier. In addition, we included LC-Orbitrap-MS based metabolomics analysis to gain a closer view on the impact of soy lecithin on functionality. In this research paper, we thus describe the impact of soy lecithin on the composition and functionality of a high-responder and a low-responder microbiota.

## 2. Materials and methods

### 2.1 Experimental design of the SHIME-model

The impact of soy lecithin on the human gut microbiota was investigated using the mucosal simulator of the human intestinal tract (M-SHIME^®^), a dynamic semi-continuous reactor-based model that mimics the different stages of the human gastrointestinal tract (ProDigest-Ghent University, Ghent, Belgium). Two independent SHIME-experiments were set up for this study, each inoculated with the faecal gut microbiota from a different human donor. In each experiment, two concentrations (0,05 m% and 0,5m%) of soy lecithin were tested alongside a control condition that received water, in parallel sets of stomach/small intestinal vessels coupled to colon vessels (Supplementary materials Figure 1). The concentrations were chosen based on what is commonly applied by the food industry and authorised in food products by the European Food Safety Authority (EFSA) and the Food and Drug Administration (FDA) [22–24]. The mucosal environment was simulated through addition of mucin coated carriers in the colon compartments [25]. During the experiment, the double-walled vessels were continuously stirred by magnetic stirrers (Prosense, Oosterhout, The Netherlands) at 200 rpm and kept at 37°C through connection to a hot water bath (Julabo, Seelbach, Germany). The pH of the colon vessels was continuously monitored using pH electrodes (Consort, Turnhout, Belgium) and controlled at 6,15 – 6,3 by semi-automatic addition of 0,5 M NaOH or 0,5 M HCl through pH controllers (Consort, Turnhout, Belgium) and pumps (ProMinent, Heidelberg, Belgium).

### 2.2 Operation of the SHIME-experiments

On the day of start-up, a faecal sample was harvested from a previously selected donor. The donors for these experiments were chosen from a pool of 10 individuals that were consulted for previous experiments [26]. Based on the response of the gut microbiota from these 10 donors to the dietary emulsifiers in terms of SCFA-production, one high and one low responder was chosen. Both donors were female, one 24 and the other 29 years old. Both had not received any antibiotic treatment in the three months prior to the faecal donation. The faecal samples were harvested in plastic lidded containers, which were rendered anaerobic by addition of Anaerogen^™^ sachets (Oxoid Ltd., Basingstoke, Hampshire, UK), and stored at 4°C for a maximum of 1 hour. A faecal slurry was then prepared as described in De Boever et al., (2000).

Before inoculation, the SHIME-model was operated on water for 24 hours. On the day of inoculation, all tubing and vessels were emptied, the stomach/small intestinal vessels were connected to pancreatic solution, containing 12,5 g/L NaHCO3 (Sigma Aldrich, St. Louis, MO), 0,9 g/L porcine pancreatin (Sigma Aldrich, St. Louis, MO) and 6 g/L Oxgall (BD, Franklin Lakes, NJ), and nutritional SHIME feed (Prodigest PD – NM002A)(composition noted in Supplementary materials Table 1), acidified to a pH of 2,0. The colon vessels were filled with 360 mL of non-acidified nutritional SHIME medium, were then inoculated with 40 mL of faecal slurry, manually flushed with 100% N_2_ gas and left to acclimatize overnight.

The next day, the pumps were turned on and a first set of samples were taken, marking the official start of the experiment (0h). The feed – pumping scheme of the SHIME model is set out in Supplementary materials Table 2. Briefly, the stomach vessels were fed 140 mL of nutritional medium 3 times a day, as well as 60 mL of pancreatic juice, the feeding times 8 hours apart. From the stomach vessels, the solution was pumped to the colon vessels and after a residence time of 16h, discarded.

After at least 5 days of stable operating conditions, which was evaluated by means of pH-stability and visual inspection of the model, the microbial community was considered adapted to the SHIME-environment and the treatment was started. The pH was considered stable when it remained within a range of 6,15 −6,3 and visually, conditions were considered stable if the contents of the colon vessels remained turbid, indicating proper bacterial growth. Treatment solutions were prepared from Barentz Unilecithin soy lecithin (UNILEC– ISL non GMO IP) in distilled water and 10x more concentrated than the intended concentrations in the vessels. Three times a day, 40 mL of treatment solution was pumped into the respective stomach/small intestine vessels. After a treatment period of 1 week, the experiment was terminated.

To maintain anaerobic conditions during the experiments, all 14 SHIME-reactors were flushed for 5 min with N_2_ daily, between 9 and 10 am, except on days where mucin beads were replaced, in which case flushing occurred after bead replacement, around noon. Luminal samples were taken on days 2 (0h), 3, 5, 7, 8, 9, 10, 11 (twice: before and 1h after the start of the emulsifier treatment), 12, 13, 14, 16 and 18 (432h) for SHIME experiment 1 and days 2 (0h), 3, 5, 7, 8, 9, (twice: before and 1h after the start of the emulsifier treatment), 10, 11, 12, 14 and 16 (384h) for SHIME experiment 2. The full sampling scheme can be found in Supplementary materials Table 3 and 4. The procedure for taking luminal samples entailed attaching a 10 mL syringe to the sampling outlet on the lids of the colon reactors of the model, rinsing the syringe and the sampling tube of the colon vessel with the content of the vessel by filling and emptying the syringe multiple times, finally taking a 10 mL aliquot and dividing it in volumes of 1 mL over 1,5 mL Eppendorf tubes and one 10 mL tube for short chain fatty acid (SCFA) analysis.

Half of the 60 mucin beads (4 clusters of 15 beads) in each of the colon vessels were replaced every 2 days (except on weekends). First, mucin beads were coated in agar-mucin as described in Van den Abbeele et al., (2012). The beads were then sown together in clusters of 15 and stored in a sterilized container at 4°C until use. In the vessels, the beads were hanged inside a polyamide net, attached to lid with a nylon wire to facilitate exchange. Beads were always replaced in the morning between 10 and 12 am in the following way. First the lids of the vessels were connected with the N_2_ flush system and the vessels would be flushed to ascertain anaerobic conditions during the whole procedure. Next, the vessels were opened one by one, 2 of the 4 clusters in the vessels were replaced by 2 fresh ones, the vessels were closed and flushed with N_2_-gas for a further 5 min. Meanwhile, the beads were centrifuged for 3 minutes at 500 *g* to collect the mucin.

### 2.3 SCFA-analysis

The SCFA - concentrations in the luminal SHIME-suspensions were determined throughout the experiment by means of diethylether extraction and capillary gas chromatography coupled to a flame ionization detector [28,29]. To this end, 1 mL aliquots were diluted 2x with 1 mL mili-Q water and SCFA were extracted by adding approximately 400 mg NaCl, 0,5 mL concentrated H_2_SO_4_, 400 μL of 2-methyl hexanoic acid internal standard and 2 mL of diethyl ether before mixing for 2 min in a rotator and centrifuging at 3000 *g* for 3 minutes. Upper layers were collected and measured using a GC-2014 capillary gas chromatograph (Shimadzu, s’ Hertogenbosch, the Netherlands), equipped with a capillary fatty acid-free EC-1000 Econo-Cap column (Alltech, Lexington, KY, US), 25 m × 0.53 mm; film thickness 1,2 μm, and coupled to a flame ionization detector and split injector. Resulting SCFA-levels were visualized in R (version 4.1.0) by use of lineplots created using ggplot2 (v3.3.3).

### 2.4 Total cell counts

To assess the impact of the emulsifiers on total cell concentrations, SYBR^®^ green staining was performed on luminal SHIME-samples after which cells were counted on an Accuri C6+ Flow cytometer from BDbiosciences Europe. Samples were analyzed every few days in batches from frozen at −20°C. Dilutions up to 10 000x were prepared in 96-well plates using 0,22 μm filtered sterile 0,01 M phosphate buffered saline (PBS) (HPO_4_^2-^/H_2_PO_4_^-^, 0,0027 M KCl and 0,137 M NaCl, pH 7,4, at 25 °C) and these were subsequently stained with SYBR^®^ green (SG) (100x concentrate SYBR^®^ Green I, Invitrogen, in 0,22 μm-filtered dimethyl sulfoxide) [30,31]. After incubation for 25 minutes, the total populations were measured immediately with the flow cytometer, which was equipped with four fluorescence detectors (530/30 nm, 585/40 nm, >670 nm and 675/25 nm), two scatter detectors and a 20 mW 488 nm laser. The flow cytometer was operated with Milli-Q (MerckMillipore, Belgium) as sheath fluid. The blue laser (488 nm) was used for the excitation of the stains and a minimum of 10 000 cells per sample were measured for accurate quantification. Settings used were an FLH-1 limit of 1000, a measurement volume of 25 μL and the measurement speed was set to fast’. Cell counts were obtained by gating the total cell populations in R (version 4.1.0) according to the Phenoflow-package (v1.1.6)[32]. Gates were verified using data from cell-free control samples (0,22 μm filter sterilized 0,01M PBS) (Supplementary Figures 1). Lineplots of total cell counts were created using ggplot2 (v3.3.3).

### 2.5 Amplicon sequencing

Luminal samples from day 2, 11, 12, 14, 18 from SHIME 1 and day 2, 9, 10, 12 and 16 from SHIME 2 and mucosal samples from day 4, 11, 14, 16 and 18 from SHIME 1 and day 4, 9, 11, 14 and 16 from SHIME 2 were selected for Illumina 16S rRNA gene amplicon sequencing. Extraction and quality verification of DNA, library preparation and 16S rRNA gene amplicon sequencing were performed as described in Miclotte et al., (2020).

Recovery of Operational Taxonomic Units (OTUs) from the amplicon data was carried out using mothur software version 1.40.5 and guidelines [33]. First, contigs were assembled, resulting in 9 815 109 sequences, and ambiguous base calls were removed. Sequences with a length of 291 or 292 nucleotides were then aligned to the silva_seed nr.123 database, trimmed between positions 2 and 13 423 [34]. After removing the sequences containing homopolymers longer than nine base pairs, 2 222 311 (99%) of the unique sequences were retained. A pre-clustering step was then performed, allowing only three differences between sequences clustered together and chimera.vsearch was used to remove chimeras, retaining 73% of the sequences. The sequences were then classified using a naïve Bayesian classifier against the Ribosomal Database Project (RDP) 16S rRNA gene training set version 16, with a cut-off of 85% for the pseudobootstrap confidence score. Sequences that were classified as Archaea, Eukaryota, Chloroplasts, unknown, or Mitochondria at the kingdom level were removed. Finally, sequences were split at the order level into taxonomic groups using the OptiClust method with a cut-off of 0.03. The data were classified at a 3% dissimilarity level into OTUs, resulting in a .shared (count table) and a .tax file (taxonomic classification).

For the entire dataset of 140 samples, 137 458 OTUs were detected in 199 genera. An OTU was here defined as a collection of sequences with a length between 291 and 292 nucleotides and with 97% or more similarity to each other in the V4 region of their 16S rRNA gene after applying hierarchical clustering.

#### Bioinformatics in R

The shared and taxonomy files resulting from the mothur pipeline were loaded into R for further processing. Absolute singletons (OTUs with only one read over all samples) were removed, resulting in 28 128 OTUs being retained [35]. Rarefaction curves were created to evaluate the sequencing depth (Supplementary Figure 2) [36]. Relative and absolute abundances of the OTUs and genera were calculated from the read counts and were explored via bar plots using ggplot2 (v3.3.3) and via principle coordinate analysis (PCoA) on the abundance based Jaccard distance matrix using the prcomp-function in the stats (v4.1.0) package. Relative abundances were calculated as percentages of the total read counts per sample. Absolute abundances (cfr. quantitative microbial profiling) were calculated by multiplying the total cell counts obtained via flow cytometry with the relative abundances of the OTUs (similar to Vandeputte et al., (2017)).

The Chao1, Shannon and Simpson diversity indices were calculated for the microbial community based on the OTU-table using the SPECIES (v1.0) package and the diversity function in the vegan (v2.5-7) package. Indices were plotted using ggplot2.

To investigate the effects of the individual constraints on the microbial community, a series of distance-based redundancy analyses (dbRDAs) was then performed on the scores obtained in the PCoA on the Jaccard distance matrix using the capscale function in the vegan (v2.5-7) package. Permutation tests were used to evaluate the significance of the models and of the explanatory variables (De Paepe et al., 2018). DbRDAs were created for both the relative and absolute abundance dataset. The global model included the factors Treatment (a concatenation of the factors Emulsifier and Emulsifier concentration) Timepoint, and Donor as explanatory variables and the absolute abundances of the genera as explanatory variables. In three subsequent dbRDAs, the constrained variance attributed to each of the factors was investigated by each time conditioning out all but one factor.

The results of the dbRDAs were plotted as Type II scaling correlation triplots showing the two first constrained canonical axes (labeled as dbRDA Dim 1/2) and the proportional constrained eigenvalues representing their contribution to the total (both constrained and unconstrained) variance. The sites were calculated as weighed sums of the scores of the response variables.

Finally, to evaluate which genera were likely differentially abundant between soy lecithin treatments and control, the DESeq2 package (v 1.32.0) was applied on the read count-table at genus level. To this end, only the read counts from the samples in the treatment period were used.

To streamline the DESeq-process, pre-filtering according to McMurdie & Holmes, (2014) was first applied on the read count-table, after which a genus-level table was created using the aggregate function (stats package v4.1.0). In the generalized linear model, the factor Timepoint, Donor, and Treatment were included. A likelihood ratio test was employed within the DESeq function on the reduced model, containing only the factors Donor and Timepoint, to test for the significance of the model. Low count genera were subjected to an empirical Bayesian correction [38]. For pairwise comparison of soy lecithin treatments versus controls, Wald tests were used after shrinkage of the Log2FoldChange (L2FC) values by means of the lfcShrink function with the adaptive shrinkage estimator from the ashr-package [39]. P-values were adjusted by means of a Benjamini-Hochberg procedure [38]. Results were visualized in volcanoplots, displaying the −log(adjusted p-value) *versus* the Log2FoldChange of each genus. Additionally, box plots were created showing the log-transformed pseudocounts extracted by the plotCounts function for each genus that showed significant differential abundance.

### 2.6 Metabolomics

Both targeted and untargeted polar metabolomics analyses of luminal SHIME-samples from days 2, 11, 12, 14 and 18 from SHIME 1 and days 2, 9, 10, 12, 16 from SHIME 2 were performed in collaboration with the Lynn Vanhaecke lab according to previously described protocols [40–42].

#### Polar metabolomics analysis

Polar metabolites were first extracted from the frozen SHIME-samples by means of ultra pure water (0,055 μS cm^-1^) obtained via a purified water system (VWR International, Merck, Germany). For this purpose, 1,5 mL SHIME-sample was first spiked with 75 μL internal standard solution (100 ng/μL L-alanine-d3 and D-valine-d8) and then centrifuged for 5 min at 13 300*g*. The supernatant was then filtered over a 0,22 μm polyvinylidene difluoride filter (Millipore, Billerica, MA, USA), diluted 1:5 using ultrapure water and transferred into an HPLC-vial [40].

Polar metabolites were then separated and detected using an ultra-high performance liquid chromatograph system coupled to a Q-Exactive^™^ mass spectrometer with heated electronspray ionisation (UHPLC-HRMS Thermo Fisher Scientific, San José, CA, USA) according to validated methods as described in (De Paepe et al., (2018) and De Spiegeleer et al., (2020). Separation was achieved using a Dionex Ultimate XRS 3000 system (Dionex, Amsterdam, The Netherlands) equipped with a Acquity UPLC HSS T3 column (1,8 μm particle size, 2,1 mm internal diameter, 150 mm length, Waters, Milford, MA, USA) kept at a constant 45 °C during analysis. Samples were injected in randomized order at a volume of 10 μL and eluted using a polar to apolar gradient formed using Ultrapure water (A) and LC/MS grade acetonitrile (Thermo Fisher Scientific, San José, CA, USA) (B), both acidified with 0,1% LC/MS grade formic acid (Thermo Fisher Scientific, San José, CA, USA). Elution occurred at a constant flow rate of 0,4 mL/min. The gradient profile comprised the following proportions of solvent A: program: 0 – 1,5 min at 98%, 1,5 – 7,0 min from 98% to 75%, 7,0 – 8,0 min from 75% to 40%, 8,0 – 12,0 min from 40% to 5%; 12,0 – 14,0 min at 5%, 14,0 – 14,1 min from 5 to 98%, followed by 4,0 min of re-equilibration [40,41].

Subsequent detection of the metabolites occurred using a Q-exactive^™^Orbitrap mass spectrometer (Thermo Fisher Scientific, San José, CA, USA), with a heated electrospray ionization source (HESI – II) operated in polarity switching mode (0/B/1 position). The system was operated under the following settings: m/z range of 53,4 - 800 Da, a maximum injection time of 70 ms, a mass resolution of 140 000 full width at half maximum and an automatic gain control of 10^6^ ions. The sheath, auxiliary and sweep gas (N_2_) flow rate for the HESI-II source were set at 50, 25 and 3 arbitrary units. The heater and capillary temperatures were set at 350 °C and 250 °C respectively, the S-lens RF-level was 50 V and the spray voltage was 4 kV [42].

Before analysis of the samples, the both ionization modes of the Q-Exactive system were calibrated using ready-to-use calibration mixtures (Thermo Fisher Scientific, San José, USA) for optimal mass accuracy. Sample analysis was also preceded by injection of a standard mixture followed by injection of a series of four quality control samples – obtained by pooling equal aliquots from all samples together – to allow system stabilization. Duplicate quality control samples were also added intermittently between series of 10 samples, to allow for correction of instrument instability and signal drift.

#### Data Processing

##### Targeted metabolomics

After data acquisition, peak identification and quantification was performed within Xcalibur 3.0 software (Thermo Fisher Scientific, San José, CA, USA). Metabolite peaks were manually identified using an in-house retention time library, the peak location in the standard mixture and the C^13^/C^12^ ratio according to CD 2002/657/EC guidelines. A minimal peak intensity threshold of 500 000 a.u. was employed. Peak areas were gathered as one table in Excel 2016 and normalized through division by peak area values for the internal standard (D-valine-d3). This base table containing area ratio’s was then plugged into R (version 4.1.0) for further principle coordinate analysis or split up per treatment versus the control condition for analysis in MetaboAnalyst 5.0 (2021).

Disease signatures present in the metabolite dataset were explored using the quantitative Enrichment Analysis application in MetaboAnalyst 5.0 (Xia Lab, McGill University, Quebec, Canada). Prior to these analyses, log transformation and Pareto scaling were performed via the website tool. The faecal library was chosen as reference database. The Statistical Analysis application on MetaboAnalyst was also employed to identify differential compounds. Compounds were considered differential when they were indicated as significantly different by the Wilcoxon Rank-sum test, the VIP value was over 1 in PLS-DA and if p(1) > 0,1 and p(corr)[1] > 0,4 in OPLS-DA for treatment with both concentrations with soy lecithin.

##### Untargeted metabolomics

Compound Discoverer 3.0 (Thermo Fisher Scientific, San José, CA, USA) was used first for automated peak extraction, peak alignment, deconvolution and noise removal with settings as given in Table 1 [41,42]. Suspect compounds were identified based on the Chemspider database (Royal Society of Chemistry).

**Table 1:** Settings utilized in Compound Discoverer for peak extraction and compound detection.

| Settings in Compound Discoverer 3.0 | Value |
| --- | --- |
| Peak intensity threshold | 500 000 a.u. |
| Specific adducts | [M+H], [M+K], [M+Na], [M-H], [M-K], [M-Na] |
| Retention Time | 0,5 - 16 |
| S/N threshold | 15 |
| Max peak shift | 0,2 min |
| Mass Tolerance | <5 ppm |
| Max peak width | 0,55 |

Peak area data obtained from Compound Discoverer was plugged into SIMCA 17 (Umetrics) for multivariate statistical analysis. To this end, first QC-normalisation and QC-filtering based on a CV < 30% were performed to eliminate instrumental signal drift and metabolites for which detection was unstable. The QC-normalisation was performed per compound, by dividing the peak area in a sample by the means of the interspersed duplicate QC-samples that follow the batch of 10 samples that sample belongs to.

In SIMCA, a Principle Component Analysis (PCA) was first generated as an exploration of overall data patterns. Next, Orthogonal Partial Least Squares Discriminant Analysis (OPLS-DA) was used to generate predictive models comparing emulsifier treatment conditions to the emulsifier free control condition. For each of these comparisons a model with a Q2 > 0,5, R2Y ≈ 1 and R2X ≈ 1 were selected, of which the significance was verified by ANOVA and Permutation tests. Model creation involved compound-filtering based on the Variable Importance of the Projection criterion: VIP >1. Compounds characteristic for the particular emulsifier treatment were then identified based on their VIP (>1), a positive Jackknife confidence interval and the S-plots (|p(1))| > 0,05 and relevance | p(corr(1))| > 0,4).

## 3. Results

### 3.1 Quality control

During the SHIME-experiments, the performance of the model was evaluated by measuring and adjusting the pH in the vessels as well as following up the produced SCFA-levels and cell concentrations. These concentrations were all at normal levels, from which we concluded that the research questions can be answered.

### 3.2 Cell counts

Treatment with both 0,05 m% and 0,5 m% of soy lecithin increased the amount of cells/mL present in the SHIME vessels compared to the control, indicating growth enhancement (Figure 1). There were no big differences between both donors in terms of response.

**Figure 1:**
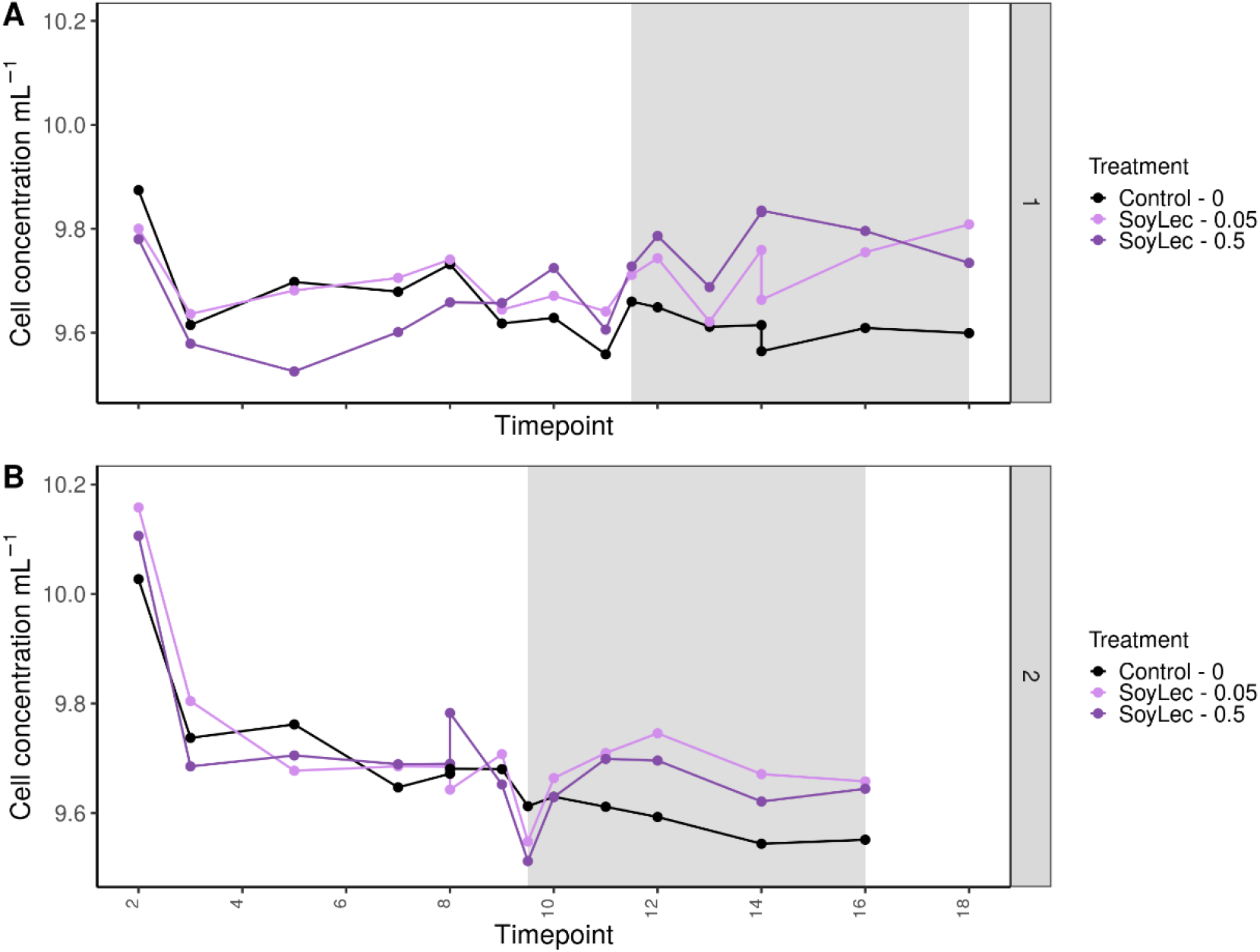
Total cell concentration (cells/mL) in luminal suspension during a 16 and 18 day M-SHIME-experiment investigating the impact of soy lecithin (0,05 m% and 0,5 m%) on the gut microbiota of two human faecal donors. Donor 1 was selected for low emulsifier sensitivity and donor 2 for high emulsifier sensitivity. The 7-day treatment period is indicated by the grey background.

### 3.3 Amplicon sequencing

To figure out which shifts in bacterial taxa were responsible for the increased total cell concentrations, the microbial composition was analyzed in luminal aliquots from day −7, 0, 1, 3 and 7 (relative to the starting day of the treatment) using 16S rRNA gene sequencing on an Illumina Miseq platform. The communities differed greatly between both donors, and their profiles were similar to those detected during the preselection. During the SHIME-experiments, the soy lecithin treatment affected different genera though, making it harder to assess whether donor susceptibility was reproduced.

PCoA and dbRDA were used to quantify the effects of the different determinants - Donor, Timepoint, Environment (mucus *versus* lumen) and Treatment – on the microbial communities. All of these factors were shown to exhibit significant effects (p < 0,005). The factor Donor was identified as the most distinguishing factor, explaining 17% of data variation for the relative abundances and 14% for the absolute abundances (Supplementary B Figure 7). The factor Timepoint explained 14% and 11% of data variation for the relative and absolute abundance data respectively. The PCoA plots indeed clearly show the distinct clustering of the microbial communities from both donors (Supplementary B Figure 3B), as well as the separation of the points at the start of the stabilisation phase *versus* those throughout the treatment phase. These plots also show a larger distance between points at the end versus the start of the treatment phase for donor 2 than for donor 1, confirming that donor 2 is the more susceptible one. The effects of the factors Environment and Treatment were less pronounced: Environment explained 7% of data variation in the relative abundance data and the soy lecithin treatment accounted for 1,17% and 5,62% of variation in the relative and absolute abundance data respectively.

The impact of soy lecithin on species richness and α-diversity was assessed by calculation of the Shannon, Simpson and Chao1 indices (Supplementary B Figures 4, 5, 6). No uniform trends were detected, though.

Given the considerable alterations in microbial structure during the stabilisation phase (Figure 3C in Supplementary B; Figure 2), the impact of soy lecithin on the microbial composition was assessed by comparison only with the datapoints just before the start of the treatment phase (Timepoint 0). Using this reasoning, the results showed that in each donor, different genera were stimulated by soy lecithin. In donor 1, treatment with soy lecithin increased the absolute abundance of the *Bacteroides*. In donor 2, rather, the *Prevotella* and Megamonas genera were stimulated. The abundance of the unclassified *Lachnospiraceae* genus was augmented for both donors (Figure 2 and 3). A more notable finding though, was that soy lecithin significantly decreased the absolute abundance of *Faecalibacterium* (*p* < 0.001) (Figure 2 and 3), a bacterium renowned for its butyrate producing capacities and for that reason considered beneficial [43,44].

**Figure 2:**
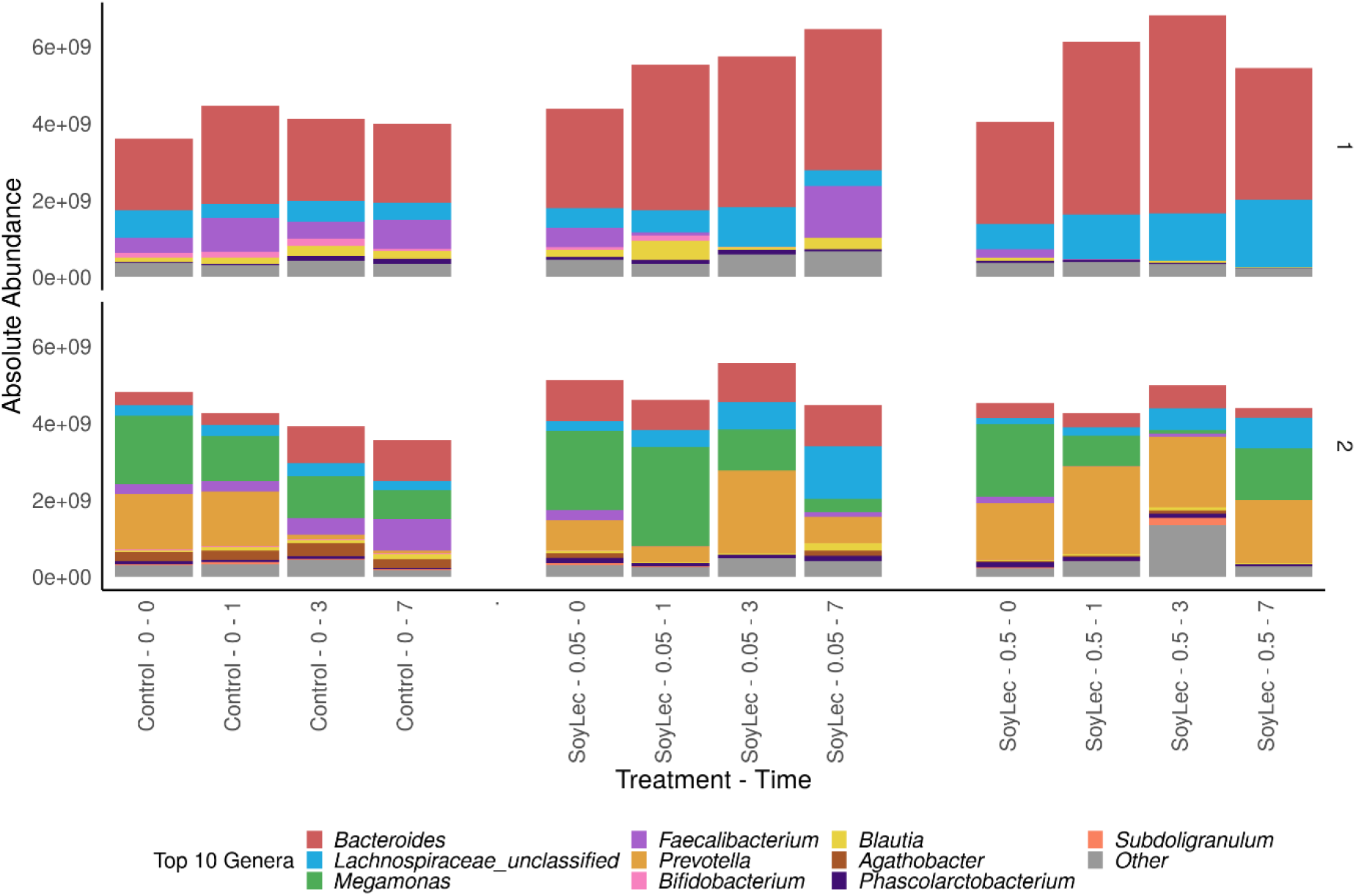
Absolute abundance plots (in cells/mL) of 10 most abundant genera derived from 16S rRNA gene amplicon sequencing, measured in luminal suspensions of a 16 and 18 day M-SHIME-experiment investigating the impact of soy lecithin (0,05 m% and 0,5 m%) on the gut microbiota from two human faecal donors. Donor 1 was selected for low emulsifier sensitivity and donor 2 for high emulsifier sensitivity. For each condition, different timepoints are indicated relative to the start of the treatment: timepoint 0 (in days).

**Figure 3:**
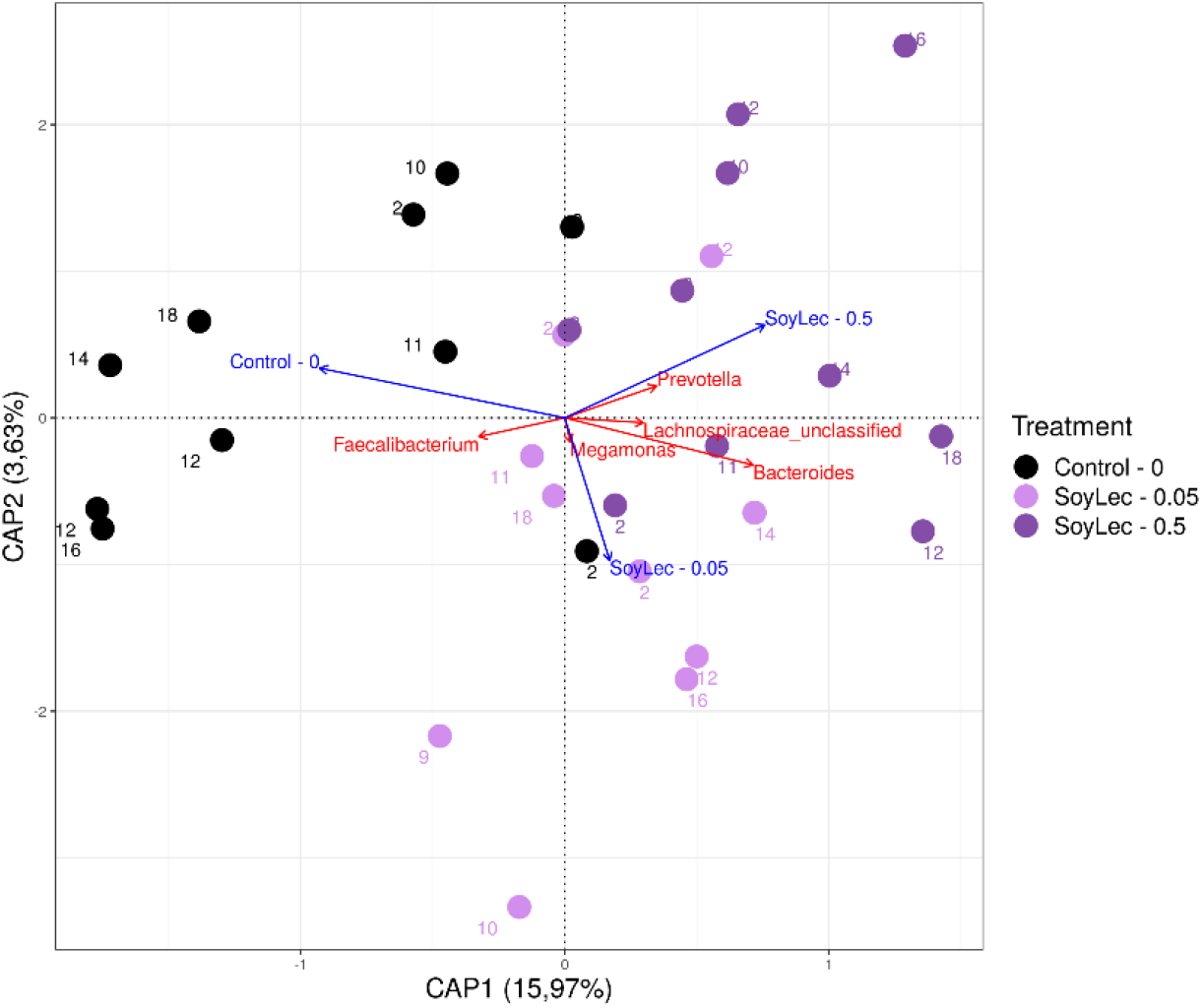
Type II scaling triplot obtained from partial distance based redundancy analysis for the absolute abundances of microbial genera in luminal suspensions from a 16 and 18 day M-SHIME-experiment investigating the impact of a 7 day treatment with soy lecithin on the gut microbiota from two human faecal donors. Absolute abundances were retrieved by use of 16S rRNA amplicon sequencing and flow cytometry. The different treatment conditions were set as explanatory variables (blue arrows) and the 5 most prevalent genera were set as response variables (red arrows). Factors Donor and Timepoint were partialled out.

To compare the differences in impact of soy lecithin on the microbial composition in the mucosal *versus* the luminal environment, we assessed the relative abundances of microbial genera detected for both microenvironments (Supplementary B Figures 8 and 9). Bargraphs and PCoA showed that the mucosal environment differed substantially from the luminal environment in both donors. The presence of the *Bacteroides* genus in the mucus was lower compared to the lumen and the relative abundance of the unclassified *Lachnospiraceae* was more notable. The mucosal environment also harboured a pronounced fraction of *Bilophila*, especially in the community from donor 2. Its relative abundance decreased, however, following treatment with soy lecithin (Supplementary B Figure 10). The relative abundance of *Lachnospiraceae* and *Faecalibacterium* was also suppressed by the treatment with soy lecithin in the mucus layer, just like in the luminal environment.

DESeq2-analysis further revealed other bacterial genera that were significantly affected by the soy lecithin treatments in the luminal and mucosal environment for both donors. In the luminal environment, 13 genera were enhanced in their growth and 23 were suppressed, whereas for the mucosal environment these 15 genera were stimulated and 11 were suppressed (Figure 4, Supplementary B Figure 11 and 12).

**Figure 4:**
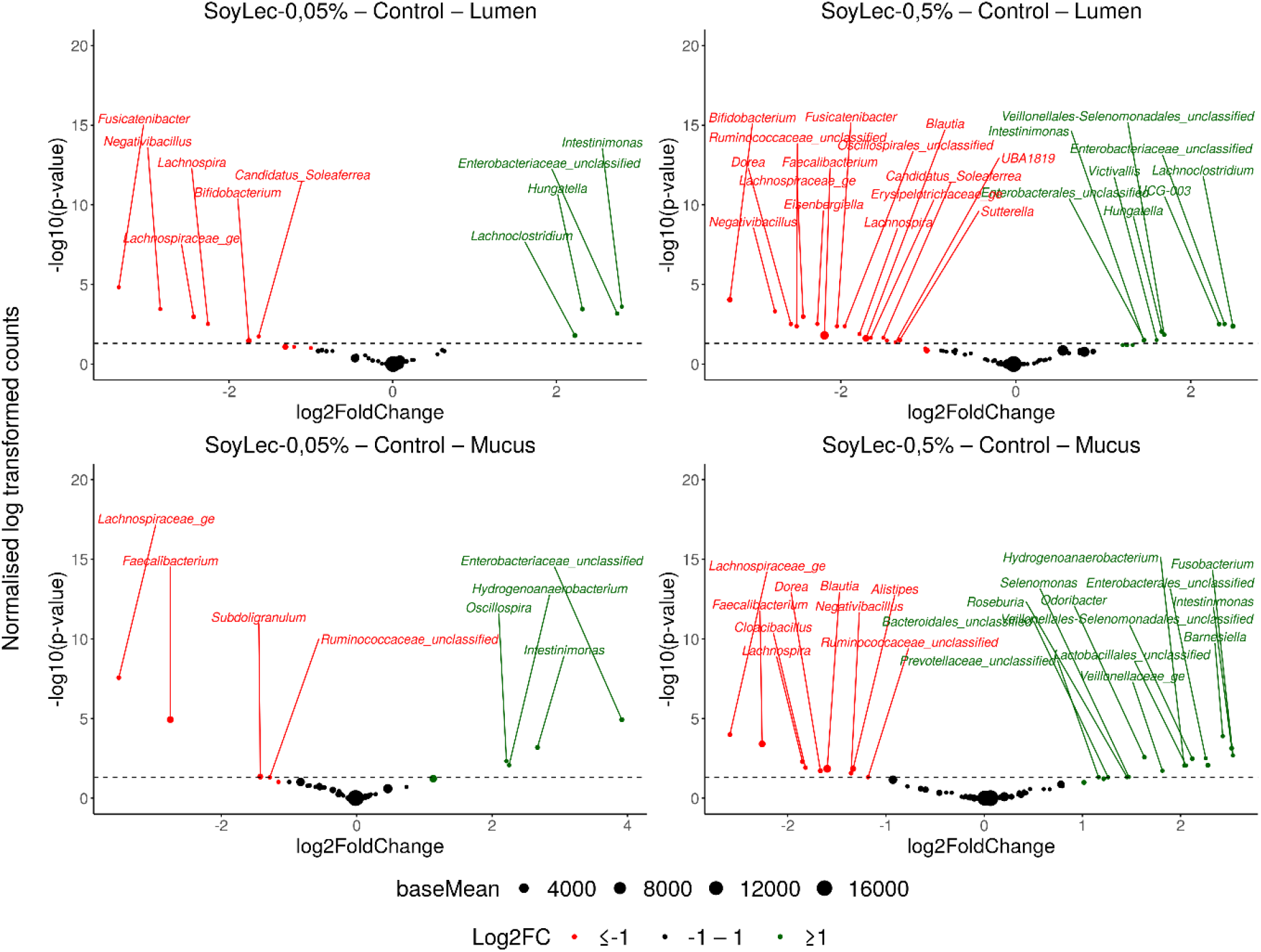
Volcano plots designating shifts to the luminal and mucosal microbial communities in response to treatment with soy lecithin (0,05 m% and 0,5 m%) during a 16 and an 18 day M-SHIME-experiment investigating the impact of a 7 day treatment with soy lecithin on the gut microbiota from two human faecal donors. Log2FoldChange (L2FC) of genus abundances for the treatments versus the controls are presented on the x-axis and the log-transformed adjusted p-value is indicated on the y-axis. Significantly in- or decreased genera are indicated in green and red respectively. The dashed line represents the significance threshold of α =0,05.

In the luminal environment, both soy lecithin concentrations significantly increased the abundance of *Hungatella* (L2FC ≥ 1,67; P_adj_ ≤ 0,009), *Intestinimonas* (L2FC ≥ 1,46; P_adj_ ≤ 0,033), *Lachnoclostridium* (L2FC ≥ 2,23; P_adj_ ≤ 0,016) and unclassified Enterobacteriaceae (L2FC ≥ 2,39; P_adj_ ≤ 0,003). Soy lecithin also significantly decreased the abundance of the genera *Bifidobacterium* (L2FC ≤ −1,76; P_adj_ ≤ 0,034), *Candidatus Soleaferrea* (L2FC ≤ −1,64; P_adj_ ≤ 0,018), *Fusicatenibacter* (L2FC ≤ −2,04; P_adj_ ≤ 0,004), *Lachnospira* (L2FC ≤ −1,79; P_adj_ ≤ 0,013), *Lachnospiraceae_ge* (L2FC ≤ −2,43; P_adj_ ≤ 0,001) *Negativibacillus* (L2FC ≤ −2,75; P_adj_ < 0,001) (Table 1in Supplementary B).

In the mucosal environment, the genera that were significantly stimulated by both soy lecithin concentrations were unclassified Enterobacteriaceae (L2FC ≥ 2,28; P_adj_ ≤ 0,009), *Hydrogenoanaerobacterium* (L2FC ≥ 2,04; P_adj_ ≤ 0,009), *Intestinimonas* (L2FC ≥ 2,52; P_adj_ < 0,001). The genera *Faecalibacterium* (L2FC ≤ −1,79; P_adj_ < 0,001), *Lachnospiraceae*_*ge* (L2FC ≤ −2,59; P_adj_ < 0,001), unclassified Ruminococcaceae (L2FC ≤ −1,18; P_adj_ ≤ 0,048) were significantly suppressed (Table 2 in Supplementary B).

**Table 2:** P-values of Wilcoxon rank sum tests (with Holm correction) comparing SCFA-value differences of treatment vessels with controls before and after the start of the treatment.

|  | Donor | Acetate | Propionate | Butyrate | Total |
| --- | --- | --- | --- | --- | --- |
| SoyLec – 0,05% | 1 | 0,69 | 0,2 | 0,34 | 0,69 |
|  | 2 | **0,03** | **0,03** | **0,03** | **0,03** |
| SoyLec – 0,5% | 1 | 0,69 | 0,34 | 1 | 0,49 |
|  | 2 | **0,03** | 0,34 | 1 | **0,02** |

### 3.4 Short chain fatty acids

Levels of SCFA were measured throughout the experiment in the luminal compartments of the SHIME to assess the impact of soy lecithin on the general functionality of the gut microbiota. In this data the emulsifier sensitivity of the donors was clear. During the treatment, total SCFA production was only significantly increased for donor 2, the most susceptible donor (Table 2 and Supplementary B Figure 13). This increase in total SCFA was mostly due to an increase in acetate levels (Table 2 and Supplementary B Figure 14). Propionate levels were slightly higher in response to soy lecithin, but only significantly for donor 2 and at the lowest soy lecithin concentration (Table 2 and Supplementary B Figure 15). Butyrate levels were decreased in donor 2 in response to the lowest soy lecithin concentration, but not the highest. In donor 1, butyrate levels from treated vessels were lower than the control curve, but not significantly (Figure 5; Table 2). These decreases in butyrate levels were significantly correlated (Corr_0,05%soy = 0,78; p_0,05%soy = 8e-5; Corr_0,5%soy = 0,674; p_0,5%soy = 0,002) with the lowered abundance of *Faecalibacterium* mentioned under 4.2.

**Figure 5:**
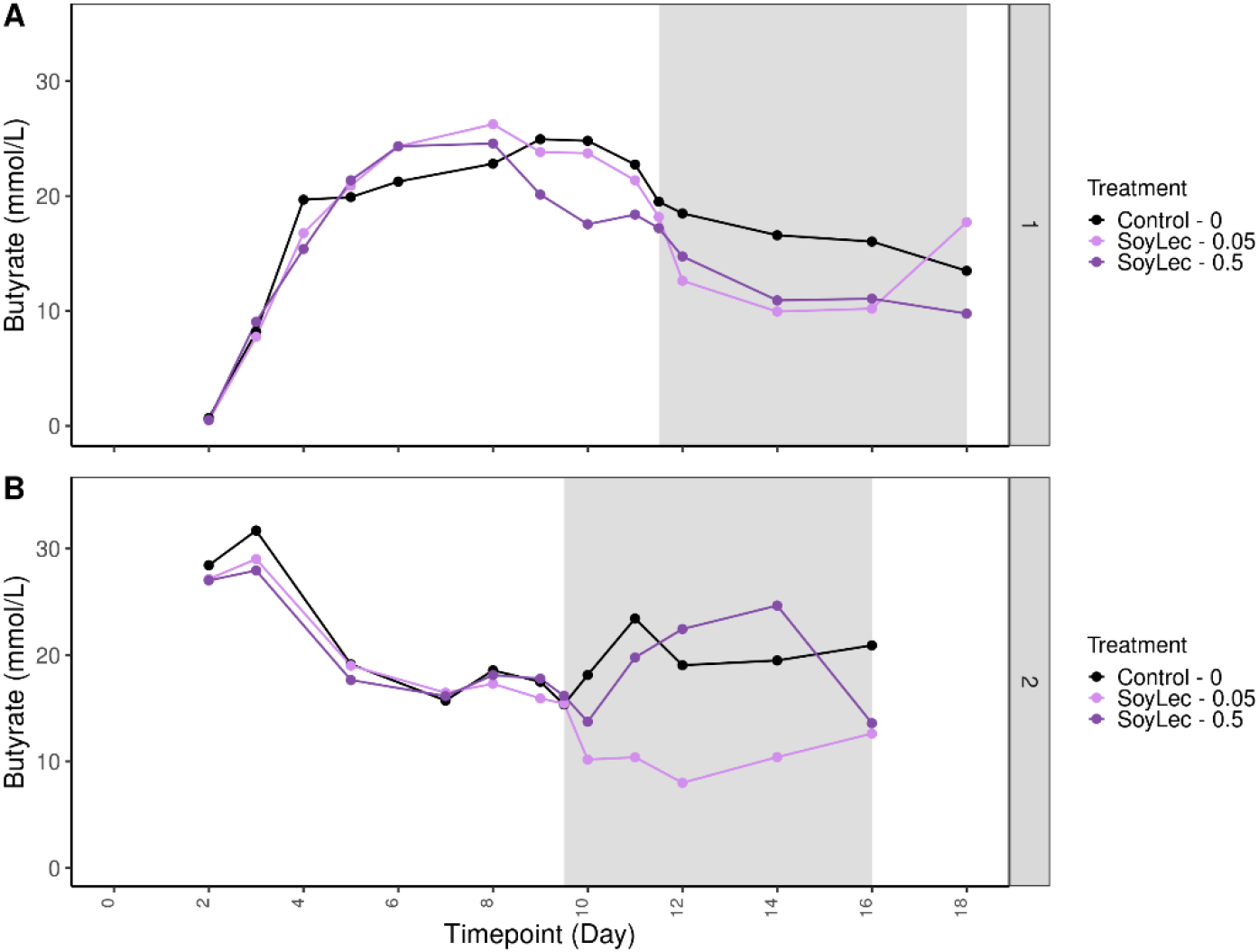
Butyrate levels (mM) in luminal suspension during a 16 and 18 day M-SHIME-experiment investigating the impact of soy lecithin (0,05 m% and 0,5 m%) on the gut microbiota of two human faecal donors. Donor 1 was selected for low emulsifier sensitivity and donor 2 for high emulsifier sensitivity. The 7-day treatment period is indicated by the grey background.

### 3.5 Metabolomics

The PCA on the extracted peak area’s from both the targeted and untargeted metabolomics data revealed slight shifts in the metabolome with the addition of soy lecithin, away from the control (Figure 6). Once more, the effect of soy lecithin was somewhat greater for donor 2 than for donor 1 (Supplementary B Figure 25). Subsequently plugging the targeted metabolomics data into the enrichment analysis tool on the MetaboAnalyst website then revealed that the metabolomic footprint resulting from soy lecithin treatment relates to those found in inflammatory bowel disease (IBD) and obesity, although this trend was not significant (Supplementary B Figures 16 and 20). The compounds of which the enrichment analysis bases these conclusions are set out in Supplementary B Figure 17-18 & 21-23. From the disease-related compounds identified for the treatment with 0,05% soy lecithin, only hypoxanthine related to an increased risk for IBD was identified as significantly different from the control (p = 0,0191). For the treatment with 0,5% soy lecithin, vaccenic acid was the only significantly different compound (p = 0,0485), which was related to an increased risk for obesity.

**Figure 6:**
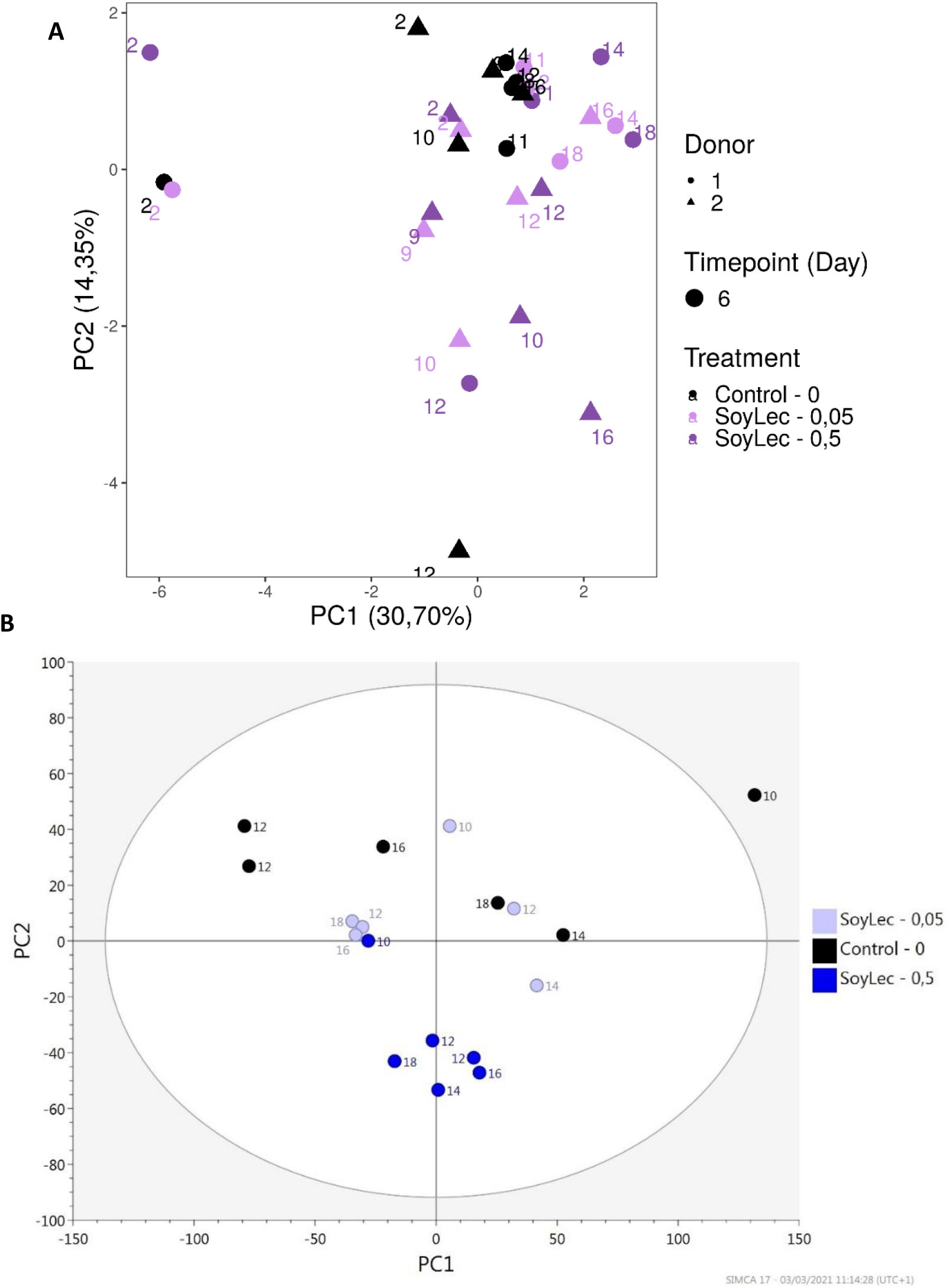
Principle coordinate analysis of targeted (A) and principle component analysis of untargeted (B) metabolomics data extracted from luminal suspensions from a 16 and an 18 day M-SHIME-experiment investigating the impact of soy lecithin (0,05 m% and 0,5 m%) on the gut microbiota from two human faecal donors.

Statistical analysis of the targeted metabolomics database further revealed significantly altered metabolites after treatment of the gut microbiota with soy lecithin. Treatment with 0,05% of soy lecithin significantly increased 5 metabolites and significantly decreased 3 (Supplementary B Figure 19). Upon treatment with 0,5% soy lecithin, 4 metabolites were significantly increased and 7 significantly decreased (Supplementary B Figure 24). Three overlapping metabolite changes were detected for both soy lecithin treatments: an increase in the levels of p-xylene and ethylbenzene and a decrease in the levels of aspartic acid (Figure 7).

**Figure 7:**
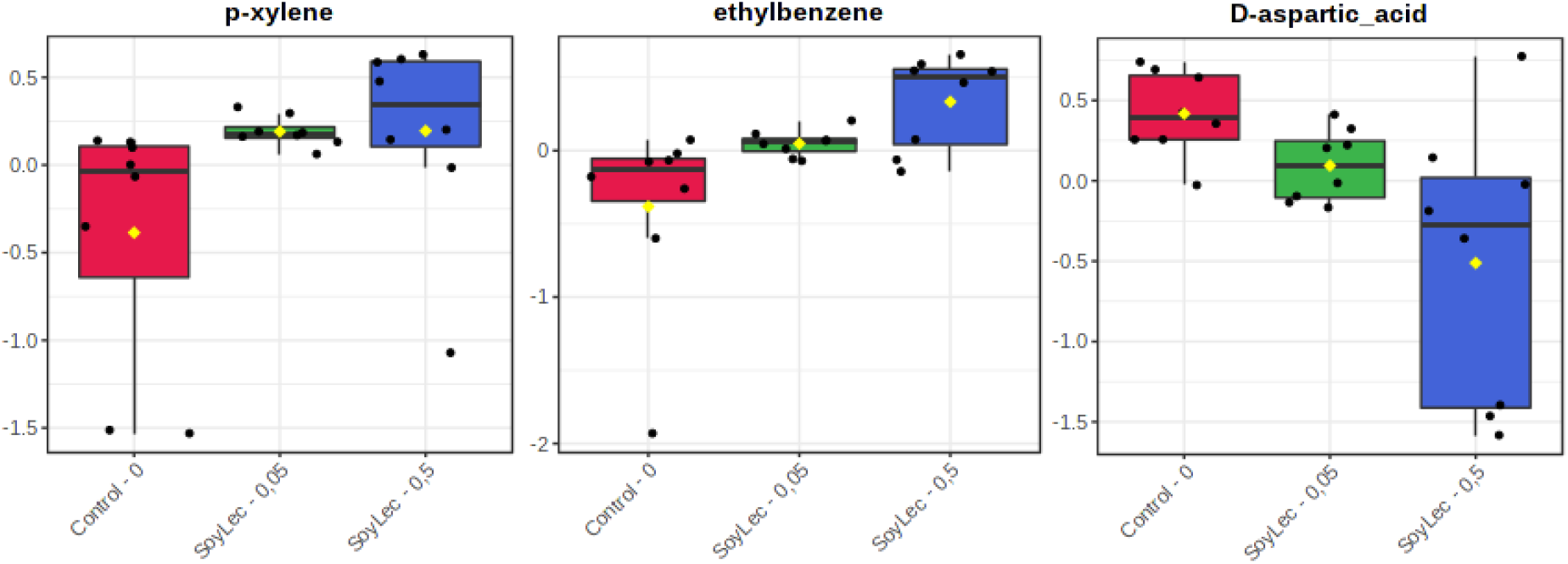
Microbial metabolites from the targeted metabolite dataset that were significantly affected by both treatments with soy lecithin (0,05% and 0,5%) in luminal suspensions from a 16 and an 18 day M-SHIME-experiment investigating the impact of soy lecithin on the composition and functionality of the gut microbiota from two human faecal donors. Compounds were considered relevant when they were indicated as significantly different by the Wilcoxon Rank-Sum test, the VIP value was over 1 in the PLS-DA and if p(1) >0,1 and p(corr)[1] > 0,4 in the OPLS-DA for treatment with both soy lecithin concentrations.

The untargeted compound extraction with Compound Discoverer (3.0) delivered a dataset of 10 953 m/z – RT pairs, of which 2977 were fully annotated using the Chemspider database. The full dataset was plugged into SIMCA for PCA and OPLS-DA modelling. OPLS-DA models were used to look for biomarkers. Compounds were considered potential biomarkers when the following criteria were met: VIP > 1, magnitude (p(1)) > 0,05 and relevance (p(corr(1))) > 0,4 (S-plot axes).

For the treatment with 0,05% soy lecithin, 4 biomarkers could be detected, whereas there were 3 for the treatment with 0,5% soy lecithin. There was one metabolite that overlapped between both treatments, which was annotated as Myxalamid A (Supplementary B Table 3). However, both soy lecithin concentrations caused those compounds to be shifted in opposite directions (Supplementary B Figure 26).

## 4. Discussion

Upfront screening of 10 individual donors upon a single dose of soy lecithin resulted in the selection of a low- and high-responding microbiota. Here, these two microbiota were studied for a prolonged time of soy lecithin exposure in the SHIME reactor, a dynamic model of the human gut. Soy lecithin altered the composition and functionality of the gut microbiota mildly and the alterations were dependent on the original architecture of the microbiota. The emulsifier sensitivity for which the donor microbiota were originally selected was preserved.

As one of the most notable findings, soy lecithin increased the bacterial cell concentration, indicative of bacterial growth, in the luminal compartment of the SHIME-system. This observation was contrary to what we observed during the 48h batch incubations of Miclotte et al (2020). For donor 1, this growth seemed to stem primarily from the *Bacteroides* and unclassified Lachnospiraceae genera, while for donor 2, unclassified *Lachnospiraceae*, *Megamonas* and *Prevotella* genera were stimulated. Other notable microbiome shifts included the suppression of the *Bifidobacterium* and *Faecalibacterium* genera and stimulation of the abundance of unclassified Enterobacteriaceae. Interestingly, soy lecithin had a selective effect on different members of the Lachnospiraceae family. On the one hand, unclassified Lachnospiraceae, *Roseburia* and *Lachnoclostridium* were stimulated, while *Blautia*, *Dorea*, *Lachnospira*, *Lachnospiraceae_ge*, *Eisenbergiella*, *Fusicatenibacter* were suppressed. This illustrates the functional diversity that exists within the Lachnospiraceae family [45].

The alterations to the microbial community by soy lecithin were different and more pronounced in this study than in our previous experimental set-up [26]. In our previous study we detected less significantly affected bacterial genera with the DESeq-algorythm and the alterations in *Megamonas*, *Prevotella*, unclassified Lachnospiraceae and *Bifidobacterium* were not significant. We did observe a slight significant decrease in *Faecalibacterium* and a slight significant increase in unclassified Enterobacteriaceae, though. To the best of our knowledge, there are only two other studies to date that have also investigated the impact of soy lecithin on the gut microbiota. The first study tested the effect of 5 days of soy and rapeseed lecithin consumption in Swiss Webster mice and found that the additive increased the abundance of *Faecalibacterium prausnitzii* non-significantly, regardless of dose or origin [46]. This finding contrasts with ours. Second, Naimi et al., (2021) have investigated the impact of 20 different dietary emulsifiers, among which soy and sunflower lecithin, on the microbial density, microbial composition, metatranscriptome, lipopolysaccharide and flagellin concentration of the human gut microbiota from one human donor in their Minibioreactor Array (MRBA) model. Although they reported few statistically significant findings regarding soy lecithin in their paper, some trends in their data were similar to ours. The microbial composition and metatransciptome were shifted slightly, and microbial richness was reduced. Also, the relative abundance of the genera of *Faecalibacterium*, *Akkermansia*, *Ruminococcaceae* and *Streptococcus* was reduced (only the latter was significant) and the genera of *Roseburia*, *Bilophila*, *Oscillospira*, as well as the families of Enterobacteriaceae and Lachnospiraceae were increased. Lipopolysaccharide-levels were increased on average by soy lecithin, which could not be said for flagellin. The trends in the abundance of *Faecalibacterium*, Rumminococcaceae, *Roseburia*, *Oscillospira*, Enterobacteriaceae and Lachnospiraceae agree with the observations in the present study. It can, thus, be concluded that soy lecithin has a mild effect on the gut microbiota, and specifically reduces the abundance of *Faecalibacterium* at least *in vitro. In vivo*, this finding could not be confirmed up till now.

Why soy lecithin stimulated the growth of certain bacterial species is unclear. Knowing that emulsification plays a role in human digestion and that phospholipids are plasma membrane components, it is possible that soy lecithin increased the availability of the supplied nutrients for microbial metabolism. On the other hand, soy lecithin may serve as an energy source itself, as it has been described that glycerol can be converted by bacteria to propionate [47]. Soy lecithin may also have altered the interactions between the different bacterial strains and in this way given one species an advantage over the other. More research into the mechanisms behind the observed quantitative alterations is required.

*Faecalibacterium* and *Bifidobacterium* are both considered beneficial bacteria. *Faecalibacterium* is known to suppress colonic inflammation, and its abundance is lowered in individuals with IBS, IBD and other gut inflammation related conditions, like apendicitis, coeliac disease, chronic diarhea and colorectal cancer [44,48,49]. *Bifidobacterium* is commonly applied as a probiotic in functional foods and is protective against constipation, colorectal cancer and gut inflammatory diseases. It also provides competition against pathogen invasion [50]. The suppression of both *Bifidobacterium* and *Faecalibacterium* in response to soy lecithin is, thus, likely an unwanted outcome.

The increased abundance of unclassified Enterobacteriaceae upon soy lecithin treatment is also potentially undesirable. The Enterobacteriaceae form a family of gram negative facultative anaerobes that contains opportunistic pathogens, like *Shigella*, *Salmonella*, *Klebsiella* and certain strains of *E. coli* [51,52]. Increased levels of Enterobacteriaceae have previously been observed in disease states, notably gut inflammation [53], obesity [54,55] and insulin resistance [55,56]. The increased presence of this family is more and more considered a signature of an unbalanced gut microbiome [57].

Functionally, SCFA analysis revealed modestly decreased levels of butyrate upon exposure to soy lecithin, compared to the control. Such lowered butyrate levels can be linked to the decreased abundance of *Faecalibacterium* genus in the lumen, which is well-known butyrate producer [43]. Interestingly, the 0,5% soy lecithin treatment for donor 2 microbiota resulted in increased butyrate levels, despite low abundances of *Faecalibacterium* in the luminal SHIME-suspension. In this case, functional redundancy in butyrate producing ability within the Lachnospireaceae or Ruminococcaceae families may explain the occupation of this functional niche. It is also possible that butyrate was produced in the mucosal compartment and diffused to the luminal suspension. As such, the *Roseburia* genus, a butyrate producer within the Lachnospiraceae family [43,58] and *Faecalibacterium* were both increased by 0,5% soy lecithin in the mucosal compartment (Figure 4).

Butyrate is considered a beneficial compound for gut health. It is the primary energy source for enterocytes and is considered anti-carcinogenic and anti-inflammatory [14,59,60]. Lower faecal butyrate levels or lower levels of butyrate-producers are frequently observed in patients with gut inflammatory diseases [49,58], and are linked with obesity-associated metabolic complications, like type II diabetes and cardiovascular problems [61–64]. Hence, lowered butyrate levels might not be desirable.

Other functional alterations were explored with metabolomics analysis. The results of the disease features analysis on the MetaboAnayst website disclosed slight shifts in the metabolome, reminiscent of those seen in IBD and obesity. Regarding IBD, several nucleotide related compounds, as well as certain amino acids were put forward as indicative. With respect to obesity, lower vaccenic acid (VA) levels were the only ones that pointed towards a more obesity prone metabolome, but this finding is somewhat dubious. Vaccenic acid is a trans-11 fatty acid generated via biohydrogenation of linoleic or linolenic acid by certain species in the gut or in the rumen of cattle. Some of these species, namely *Faecalibacterium* and *Bifidobacterium* [65,66], were suppressed in this study, providing an explanation for the lower VA-levels observed here. *In vivo*, VA is easily absorbed in the human gut and converted to conjugated linoleic acid in the liver [67] and through this conversion, consumption of VA has been related to favourable health outcomes [67,68]. However, at present, no human studies have linked VA-consumption or – production in the intestine to any form of weight loss. Moreover, it is unclear whether the gut microbiota of obese individuals displays an altered level of biohydrogenation or net vaccenic acid production in the colon. In conclusion, it is too early to start linking VA-concentrations in SHIME suspensions, or even faeces, to risk of obesity, since the mechanism of this relation is unknown.

Other altered microbial metabolites that were found in this study included p-xylene, ethylbenzene and D-aspartic acid of which the first two were significantly more present in the luminal SHIME-suspensions exposed to soy lecithin, while D-aspartic acid was detected in lower amounts in the emulsifier treated vessels. P-xylene and ethylbenzene are xenobiotics and incremented levels in the luminal SHIME-suspension suggest a lowered potential for xenobiotics degradation [69].

Before the findings observed in this study can be extrapolated to an *in vivo* situation, several questions will need to be answered. First, it is still unclear whether soy lecithin will exhibit the same effects on the gut microbiota when consumed in a food matrix. Second, several components of lecithin are digestible by human enzymes and the subsequent metabolites absorbable. It thus needs to be clarified how much of the ingested soy lecithin reaches the colon and in what form. Last, despite the fact that we selected two donors with very different emulsifier sensitivity, these long-term exposure experiments should be extended to more individuals to better capture and understand interindividual variability.

Whether soy lecithin must now be considered a harmful compound is still an open discussion with several considerations. First of all, soy lecithin and all other lecithins are comprised primarily of phospholipids [20]. These phospholipids are naturally present in all cell membranes, from bacteria to plant cells or human cells [70]. Phosphatidylcholine, a principal phospholipid in soy lecithin also overlies the gut epithelium already, since it is excreted in the intestine [71,72], so it is unclear why extra consumption of that compound via soy lecithin would be harmful. Phospholipids are also naturally present in many food products, mostly in the form of phosphatidylcholine. They are found especially in meat (0,5 – 5 g/100g), fish (0,2 – 2,5 g/100g), milk and milk product (0,03 – 0,2 g/100g), cereals (0,05 – 1,2 g/100g), oily seeds (up to 1g/100g) and legumes (2 g/100g in soybeans) [73]. Next, soy lecithin is being sold as a nutritional supplement for cardiovascular health, since it is supposed to alleviate high blood cholesterol through inhibition of cholesterol absorption [74]. However the literature isn’t entirely convincing, since too little randomised controlled trials exist. Last, phospholipids can be digested by human digestive phospholipases and its constituents - inositol, choline, serine, ethanolamine, fatty acids, glycerol – can be absorbed (Ehehalt et al., 2010; Mortensen et al., 2017; Salhi et al., 2020), so it is unclear how much of the originally ingested soy lecithin reaches the colon.

Literature has described that one of the constituents of phosphatidylcholine, namely choline, can be a cause for concern. On the one hand, choline is an essential nutrient, required for maintenance of structural integrity of cell membranes, cholinergic neurotransmission and donation of methyl groups in a few biosynthetic reactions [76,77]. However, it is also easily metabolized by the gut microbiota to trimethyl amine (TMA) which is further converted by the liver to trimethyl amine oxide (TMAO) [78,79]. TMAO is renowned risk factor for oxidative stress, a trait frequently observed in metabolic diseases [80]. TMA was, however, not detected in our analysis. Also glycerol, the central constituent of phospholipids, can be converted by the gut microbiota to on the one hand propionate, but on the other hand also to acrolein, which is toxic to DNA, proteins and signalling pathways [81]. Hence, depending on what soy lecithin metabolites are produced by the gut microbiota, possible health consequences may vary.

In summary, we have shown that soy lecithin has a mild impact on the gut microbiota from two human donors with high and low emulsifier sensitivity. Other studies have shown alterations of comparable magnitude. Some of the observed alterations do not paint a beneficial picture: the abundance of beneficial *Faecalibacterium* and *Bifidobacterium* were decreased and the abundance of Enterobacteriaceae was increased. Future research will have to clarify 1) whether and at what consumption level these observations take place *in vivo* and 2) if the effects are sufficient to contribute to the conditions that the consumption of processed food is being related to, namely metabolic syndrome.

## 5. Competing interests

The authors declare that they have no competing interests.

## 6. Acknowledgements

We thank prof. Lynn Vanhaecke and Ellen De Paepe for assisting with the metabolomics analysis and Prof. Jeroen Raes for the 16S rRNA amplicon sequencing. We also thank Jo Devrieze for providing feedback to our manuscript before submission.

## 7. Funding

This work was supported by a scholarship from the UGent special research fund (BOF17/DOC/312-file number: 01D31217), a UGent research grant (BOF17-GOA-032) and the FWO-EOS grant MiQuant.

## Supplementary materials

**Figure 1:**
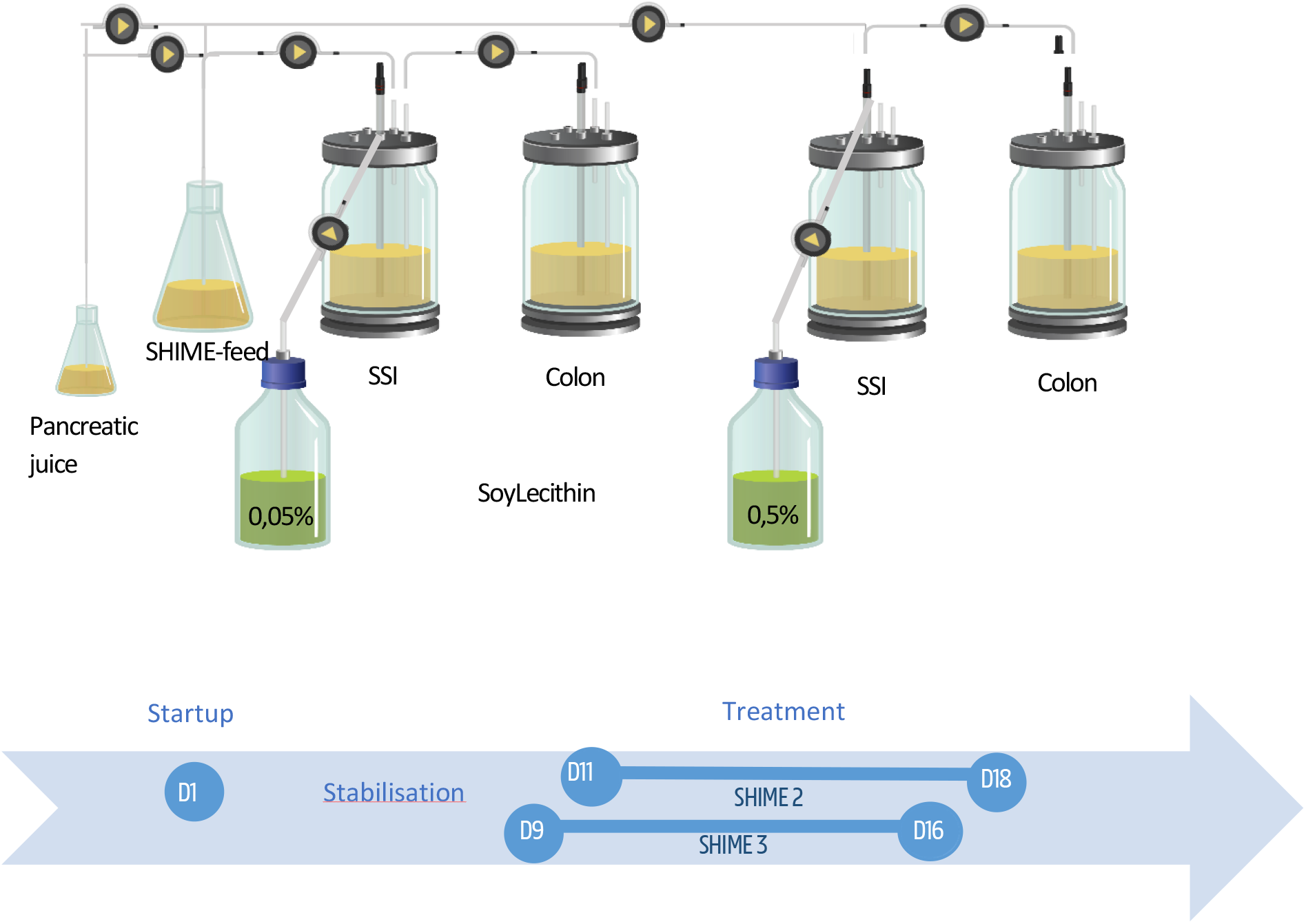
Schematic representation of an experiment with the Simulator of the Human Intestinal Microbial Ecosystem (SHIME) investigating the impact of 0,05 m% and 0,5 m% soy lecithin on the composition and functionality of the gut microbiota from 2 human faecal donors.

**Table 1:**
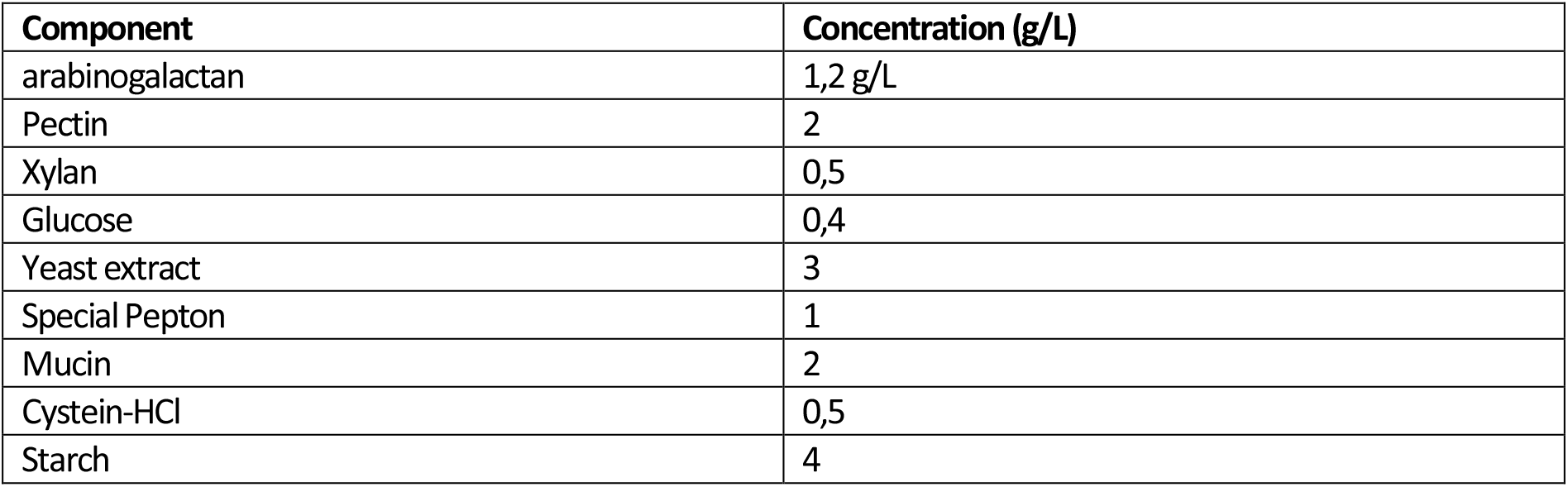
Chemical composition of nutritional SHIME-feed.

**Table 2:** Time table of pumping action between different SHIME-compartments.

| Feed =><br>Stomach/small<br>intestine | Pancreatic juice =><br>Stomach/small<br>intestine | Emulsifier =><br>Stomach/small<br>intestine | Stomach/small<br>intestine => Colon | Colon => Waste |
| --- | --- | --- | --- | --- |
| 9u -> 9u30 | 10u -> 9u15 | 9u50 -> 10u | 12u30 -> 14u30 | 13u30 -> 15u30 |
| 17u -> 17u30 | 18u -> 17u15 | 18u50 -> 19u | 20u30 -> 22u30 | 21u30 -> 23u30 |
| 1 -> 1u30 | 2 -> 2u15 | 2u50 -> 3u | 4u30 -> 6u30 | 5u30 -> 7u30 |

**Table 3:**
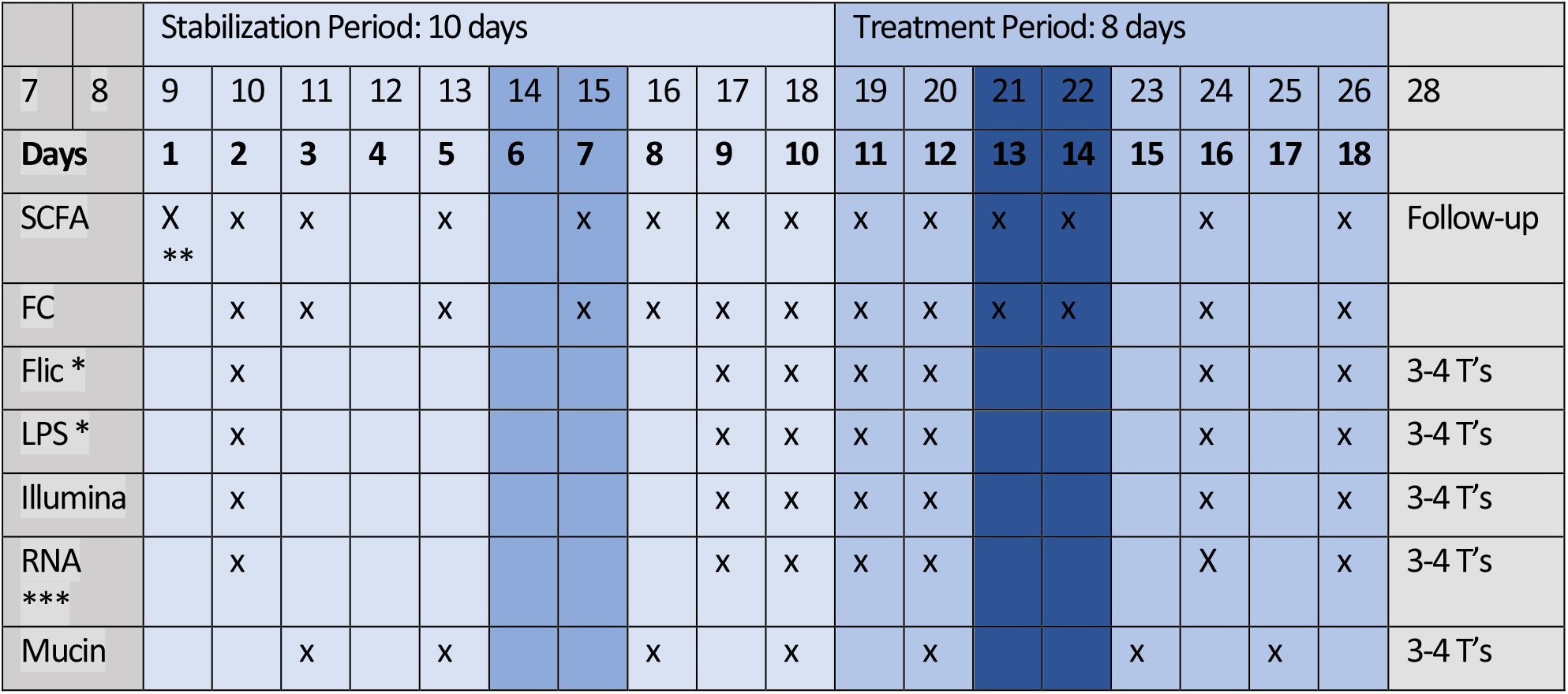
Table 3: Sampling scheme for SHIME 1 with donor 1.

**Table 4:**
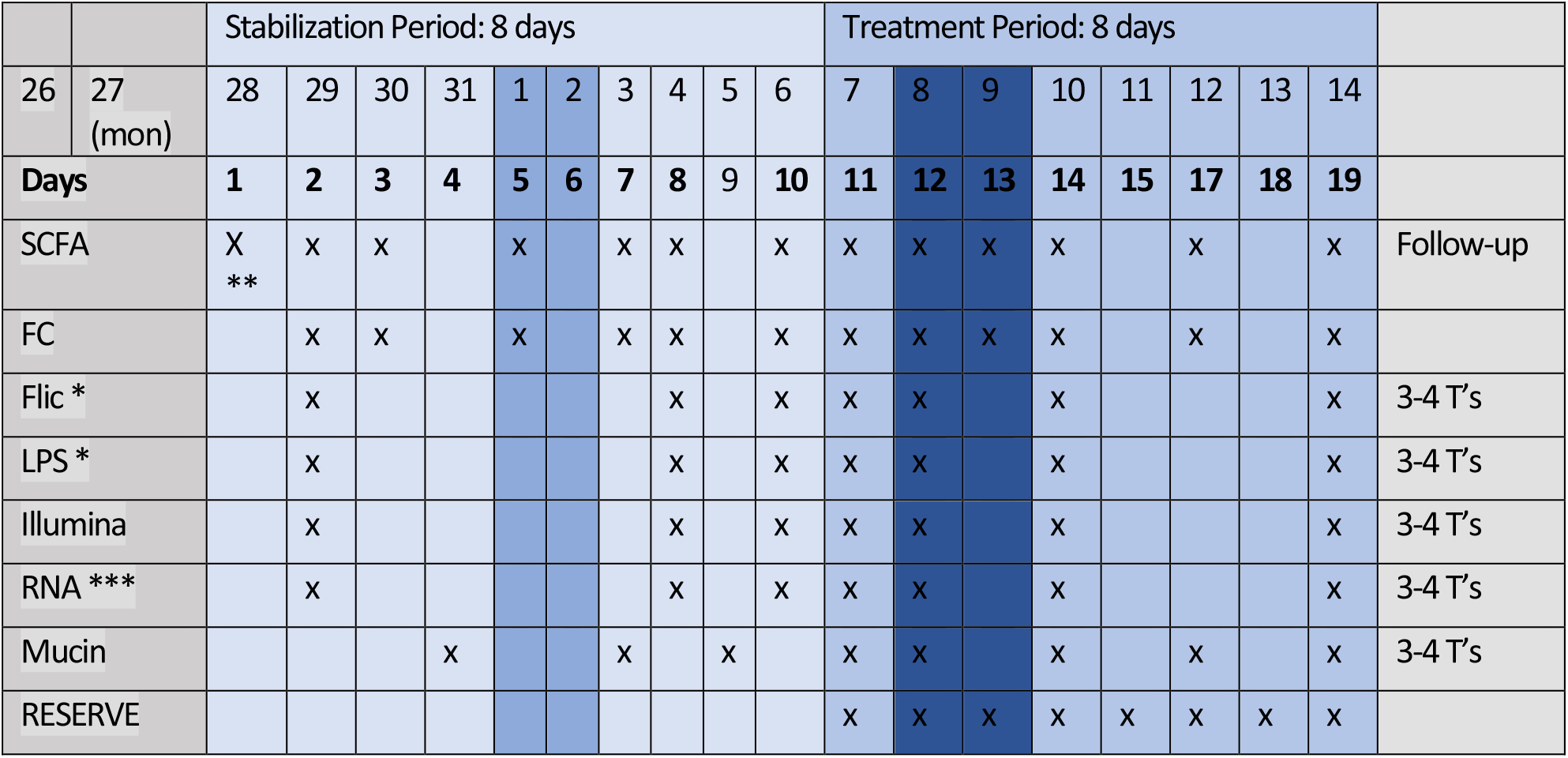
Sampling scheme for SHIME 2 with donor 2.

## Notes

### Competing Interest Statement

The authors have declared no competing interest.

