## Supplementary Figures for "Long term exposure of human gut microbiota with high and low emulsifier sensitivity to soy lecithin in M-SHIME model"

#### 1. Cell counts

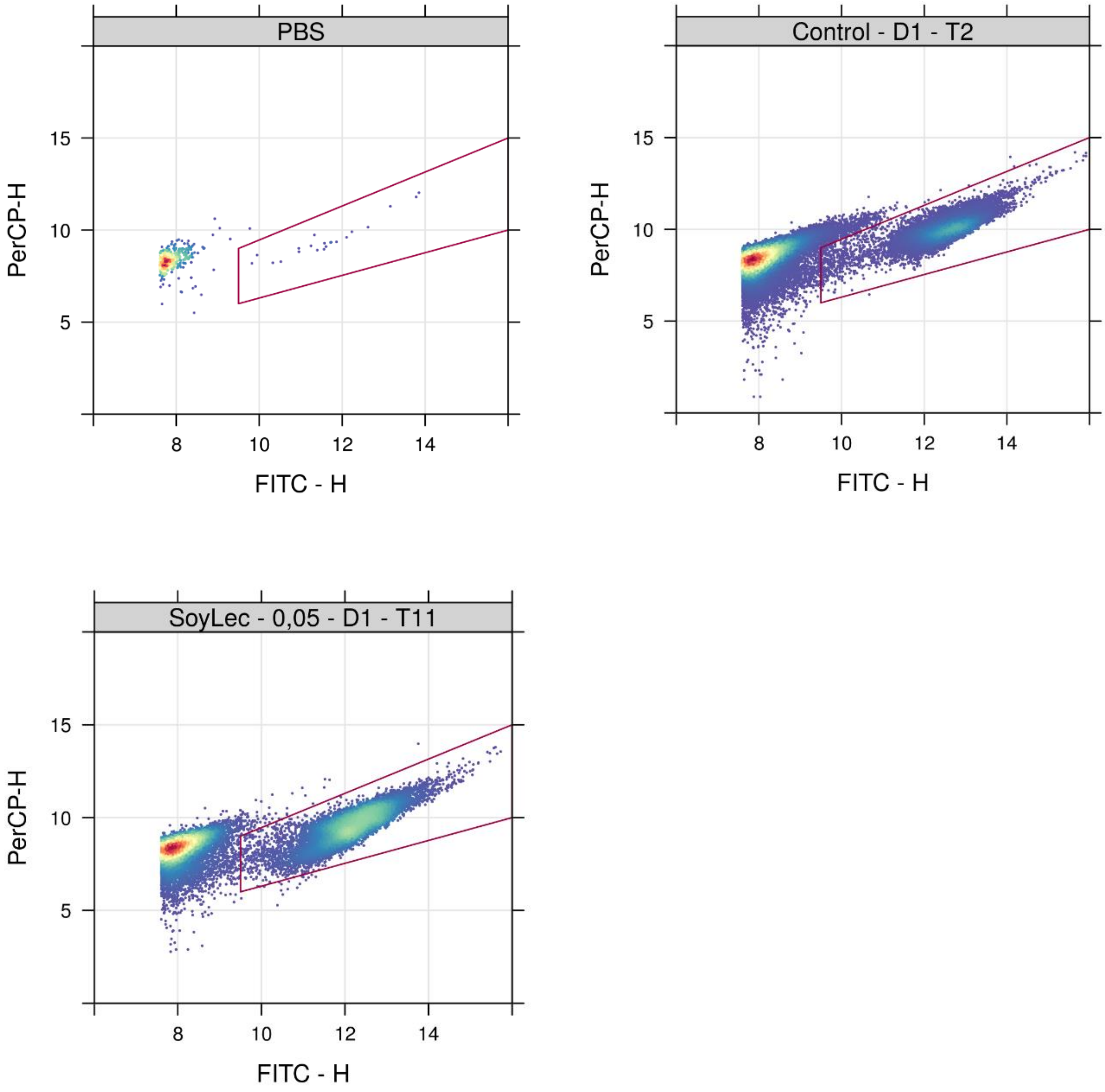


*Figure 1: Density plots of cell counts obtained through SYBR green staining and flow cytometry of several random samples from luminal suspensions from a 16 and 18 day SHIME experiment investigating the impact of soy lecithin (0,05 m% and 0,5 m%) on the gut microbiota of two human faecal donors with a 7 day treatment period. The red lines indicate the gating utilized to obtain total cell concentrations in R (version 4.1.0).*

#### 2. Amplicon sequencing

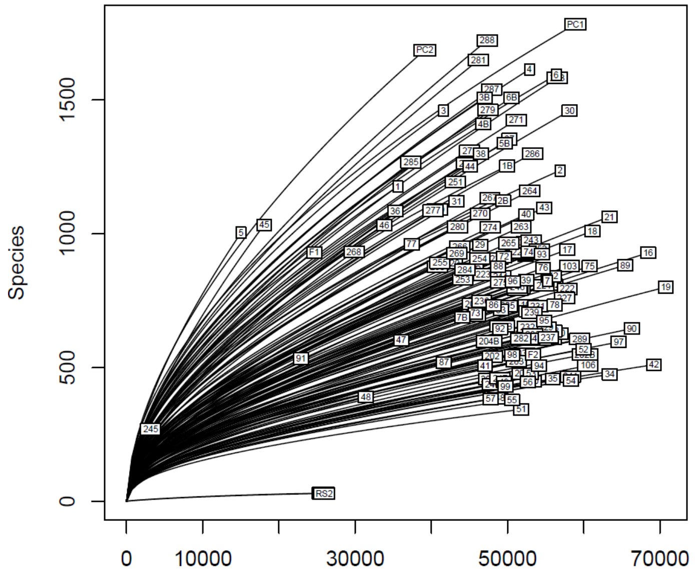


*Figure 2: Rarefaction curve of 16S rRNA gene amplicon sequencing data from in vitro M-SHIME experiments investigating the impact of soy lecithin (0,05 m% and 0,5 m%) on the gut microbiota from two human faecal donors. The numbers on the curves represent sample names.*

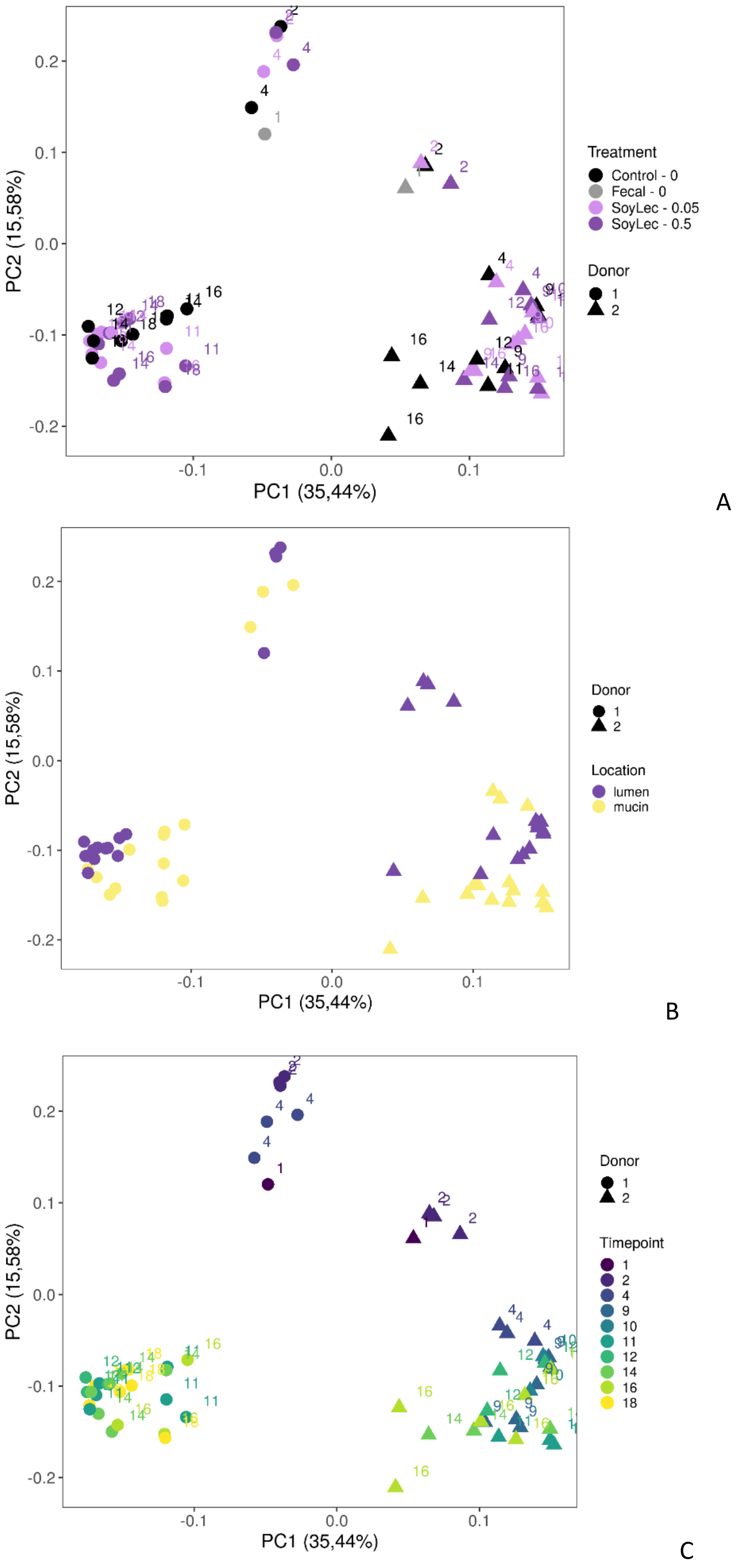


*Figure 3: Principle coordinate analysis (PCoA) of 16S rRNA gene amplicon sequencing data extracted from luminal suspensions collected during a 16 and 18 day SHIME experiment investigating the impact of soy lecithin (0,05 m% and 0,5 m%) on the gut microbiota from two human faecal donors. A: coloured by treatment; B: coloured by environment; C: coloured by timepoint.*

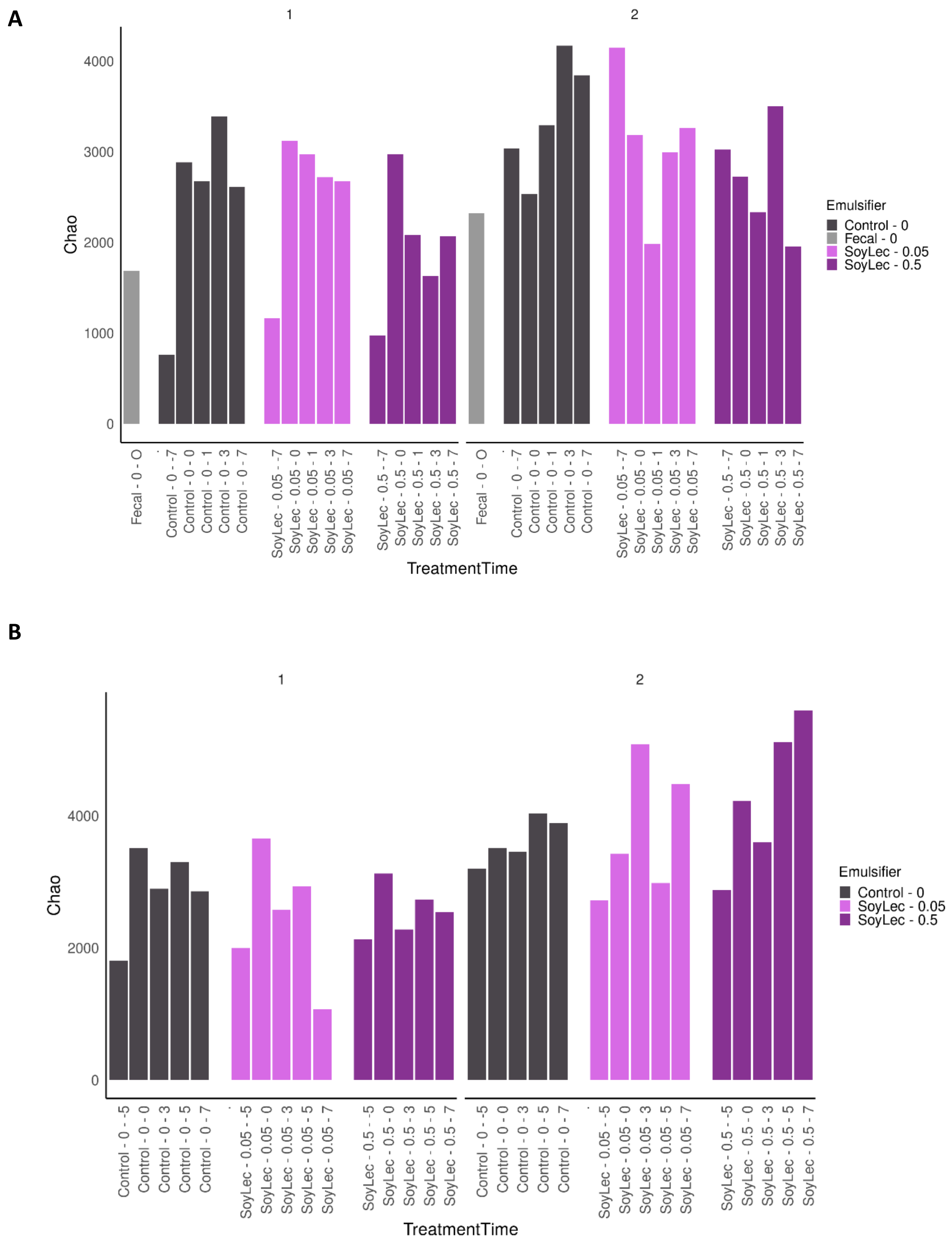


*Figure 4: Chao richness of the simulated gut microbiota in luminal suspensions from a 16 and 18 day SHIME-experiment investigating the impact of 0,05 m% (A) and 0,5 m% (B) of soy lecithin on the gut microbiota from two human faecal donors.*

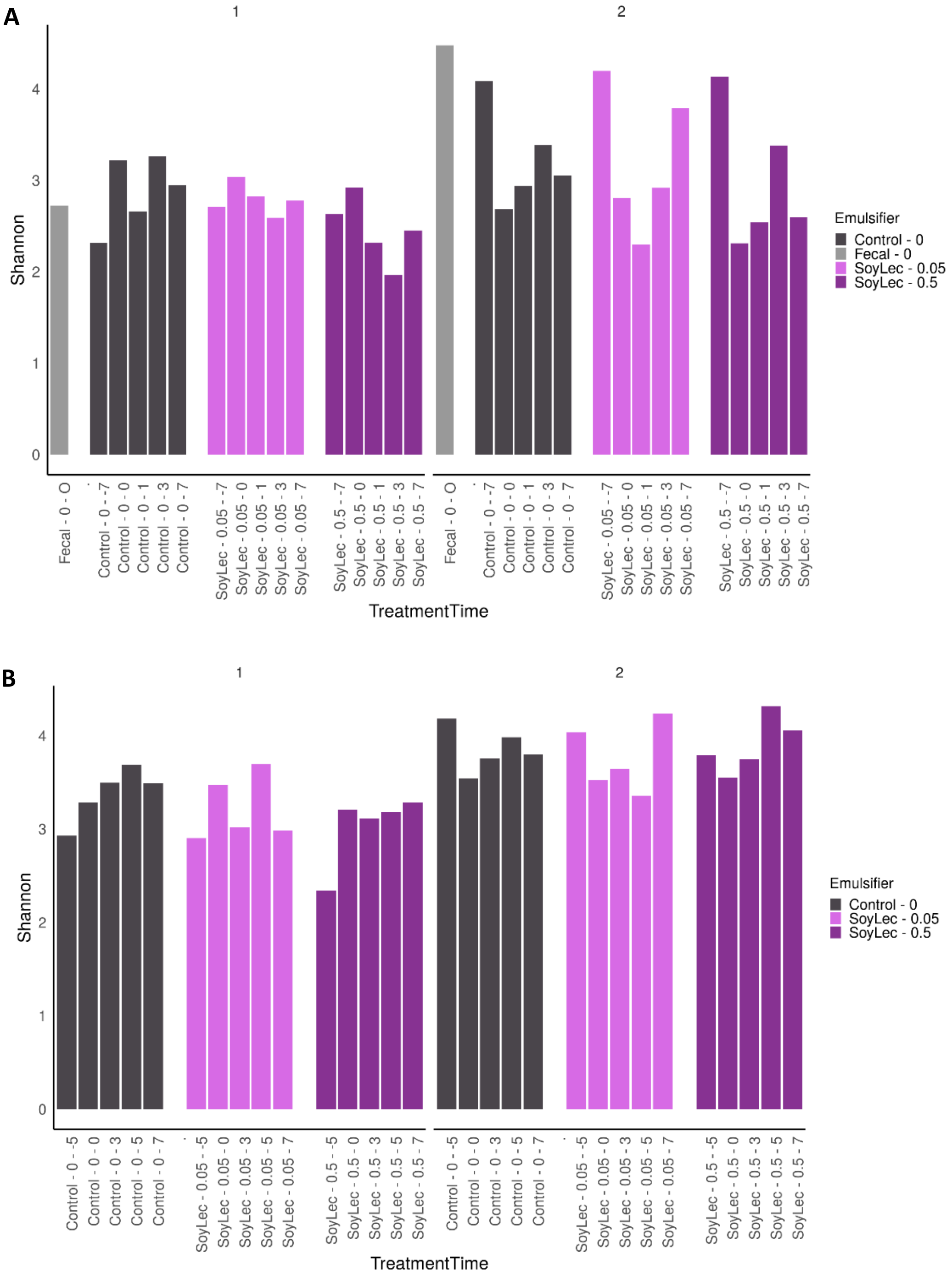


*Figure 5: Shannon index for the simulated gut microbiota in mucosal compartments of a 16 and 18 day SHIME-experiment investigating the impact of 0,05 m% (A) and 0,5 m% (B) of soy lecithin on the gut microbiota from two human faecal donors.*

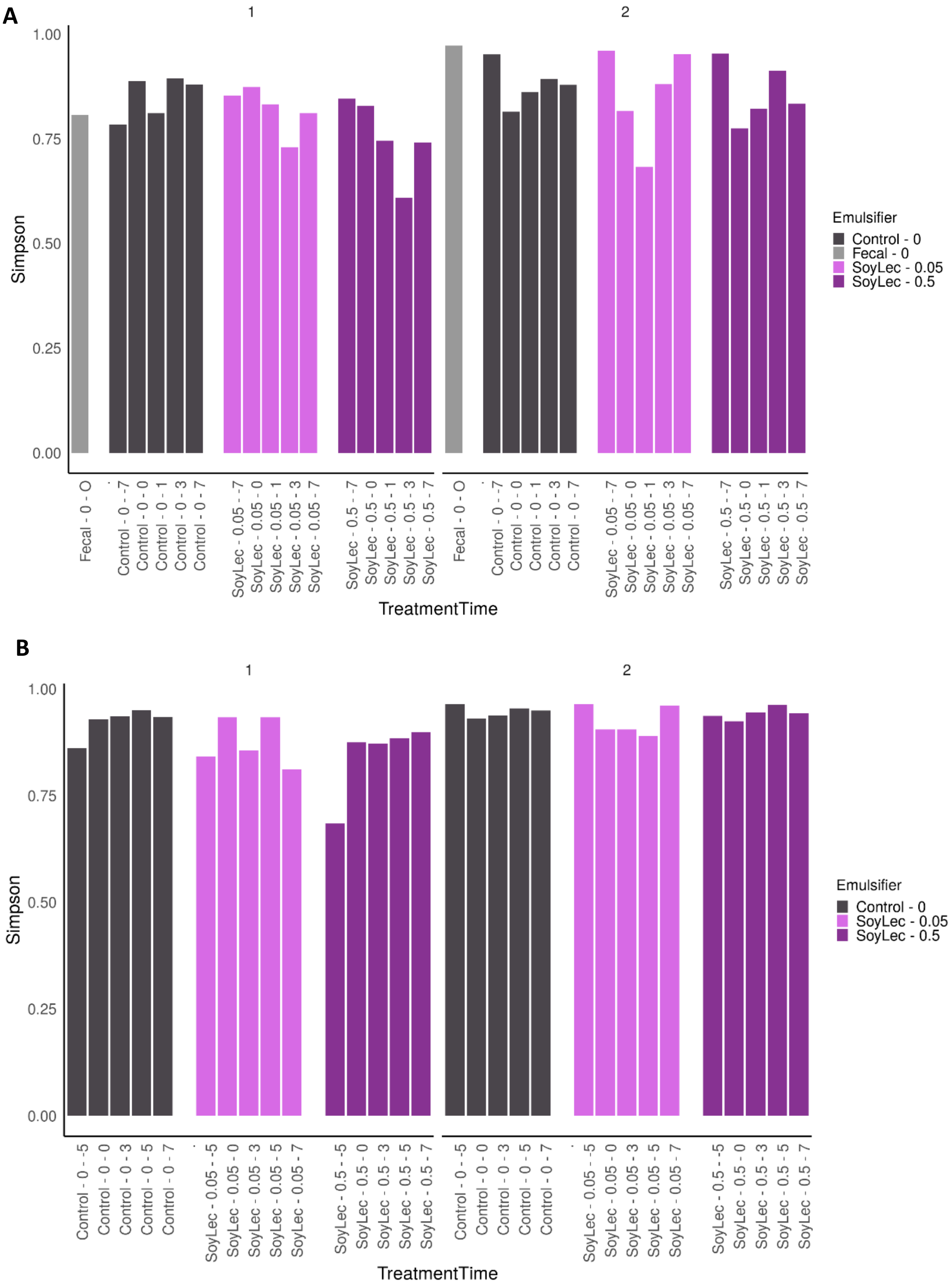


*Figure 6: Simpson index for the simulated gut microbiota in mucosal compartments of a 16 and 18 day SHIME-experiment investigating the impact of 0,05 m% (A) and 0,5 m% (B) of soy lecithin on the gut microbiota from two human faecal donors.*

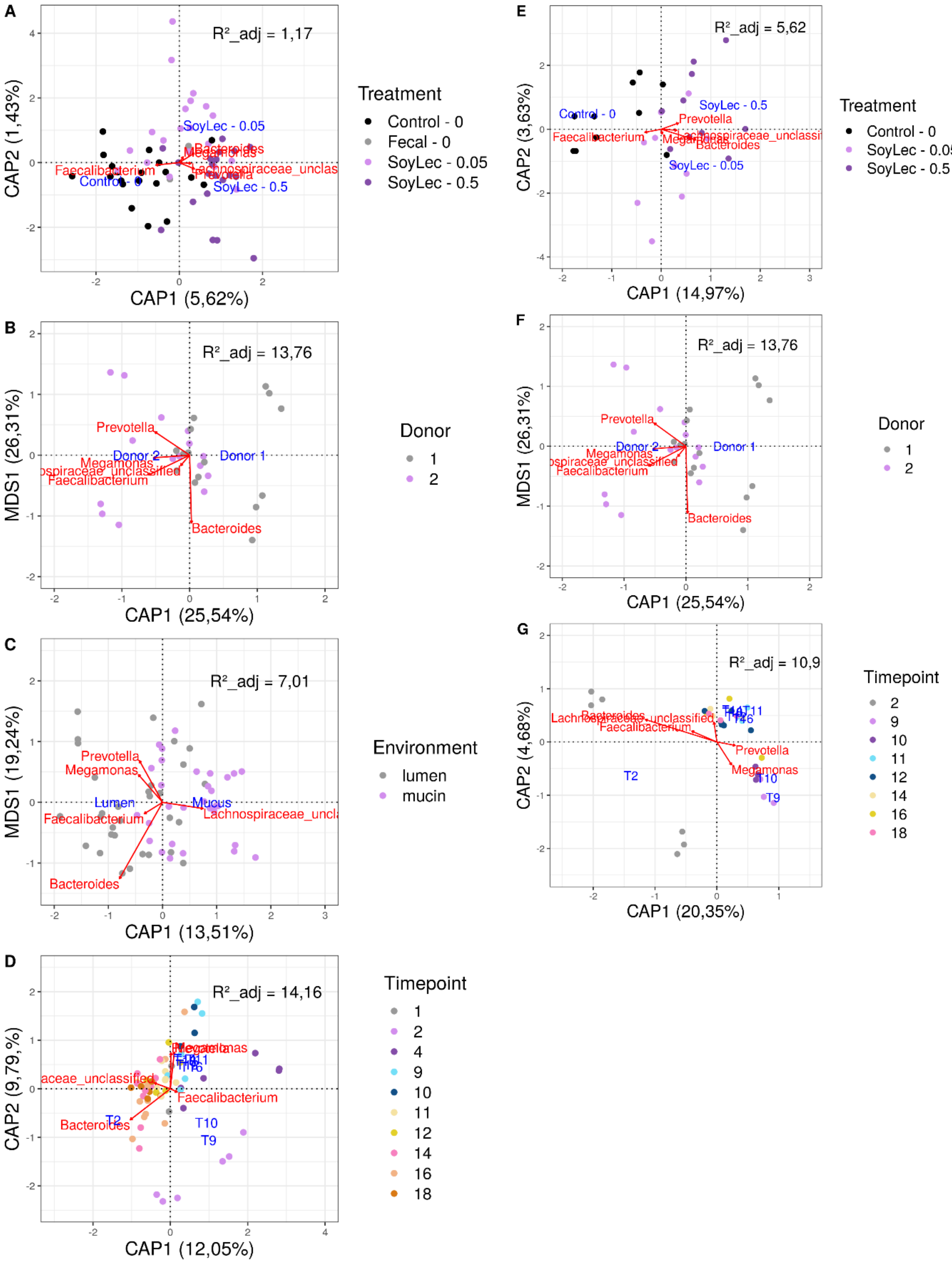


*Figure 7: Type II scaling triplot obtained from partial distance based redundancy analysis for the abundances of microbial genera in samples from a 16 and 18 day SHIME-experiment investigating the impact of a 7 day treatment with TWEEN80 and rhamnolipids (0,05 m% and 0,5 m%) on the gut microbiota from two human faecal donors. Abundances were retrieved by use of 16S rRNA amplicon sequencing. Plots A, B, C and D visualize the variation comprised by the factors Treatment, Donor, Location and Timepoint in the relative abundance data. Plots E, F and G visualize the variation explained by factors Treatment, Donor and Timepoint in the absolute abundance data. In each plot the factor levels were set as explanatory variables (blue arrows) and the 5 most prevalent genera were set as response variables (red arrows).*

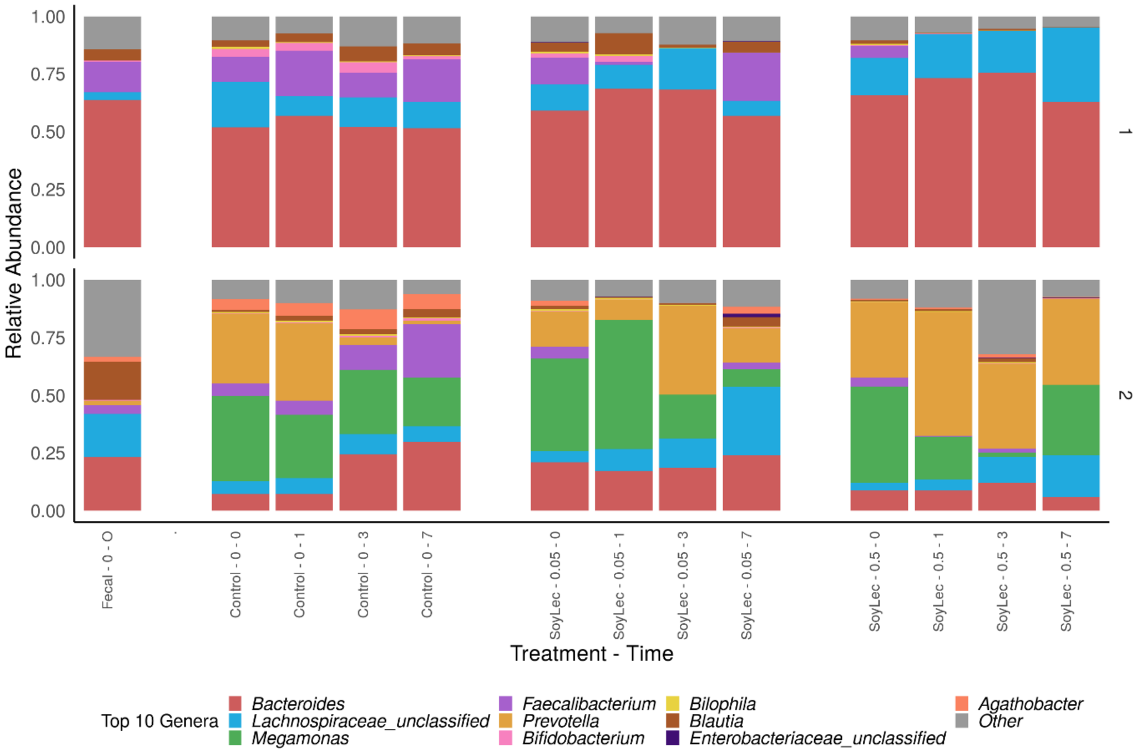


*Figure 8: Relative abundances of the 10 most abundant genera derived from 16S rRNA gene amplicon sequencing, measured in luminal suspensions of a 16 and 18 day SHIME-experiment investigating the impact of soy lecithin on the gut microbiota from two human faecal donors. For each condition, different timepoints are indicated relative to the start of the treatment: timepoint 0 (in days).*

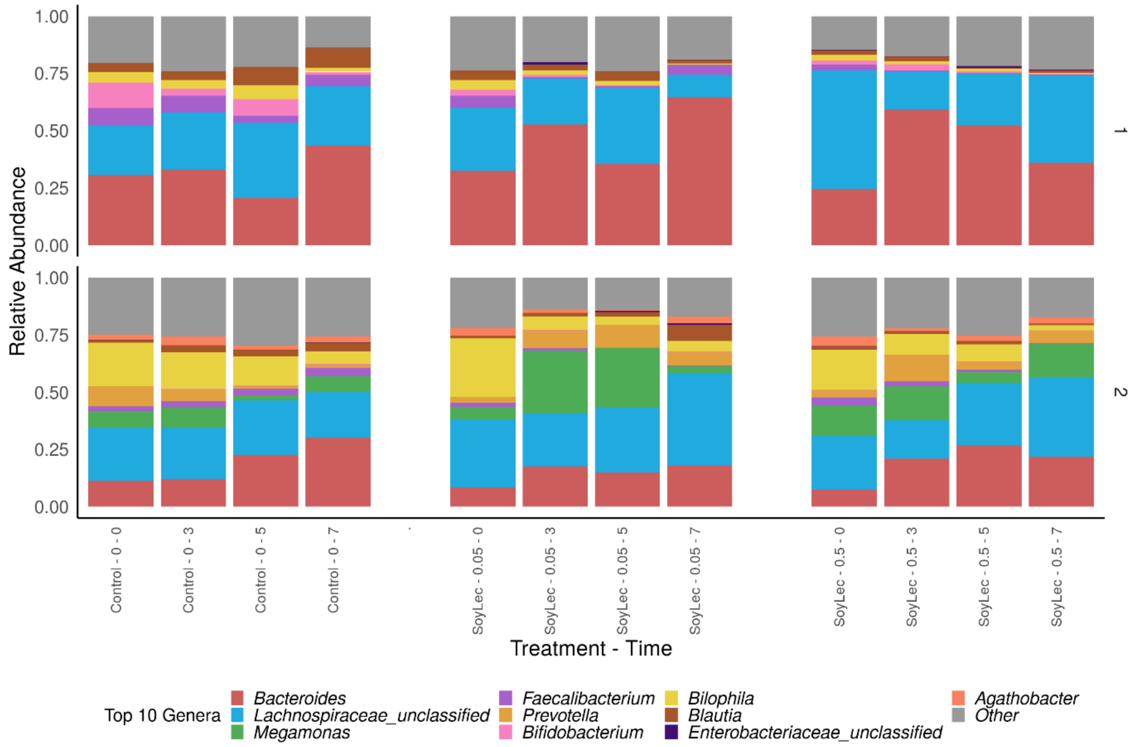


*Figure 9: Relative abundances of the 10 most abundant genera derived from 16S rRNA gene amplicon sequencing, measured in the mucosal compartments of a 16 and 18 day SHIME-experiment investigating the impact of soy lecithin on the gut microbiota from two human faecal donors. For each condition, different timepoints are indicated relative to the start of the treatment: timepoint 0 (in days).*

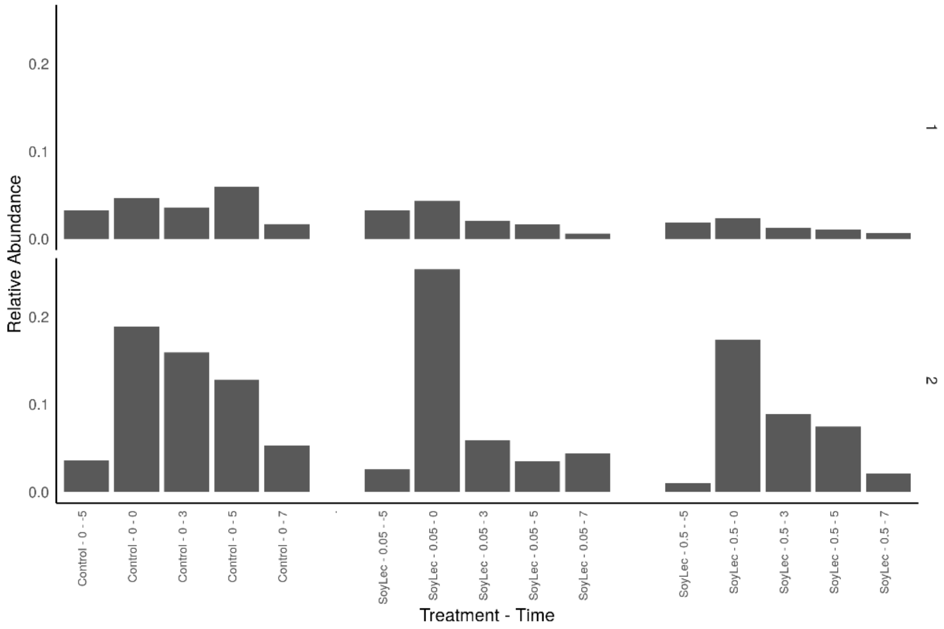


*Figure 10: Relative abundance, obtained via 16S rRNA Illumina amplicon sequencing, of Bilophila mucosal compartments of a 16 and 18 day SHIME-experiment investigating the impact of soy lecithin on the gut microbiota from two human faecal donors. For each condition, different timepoints are indicated relative to the start of the treatment: timepoint 0 (in days).*

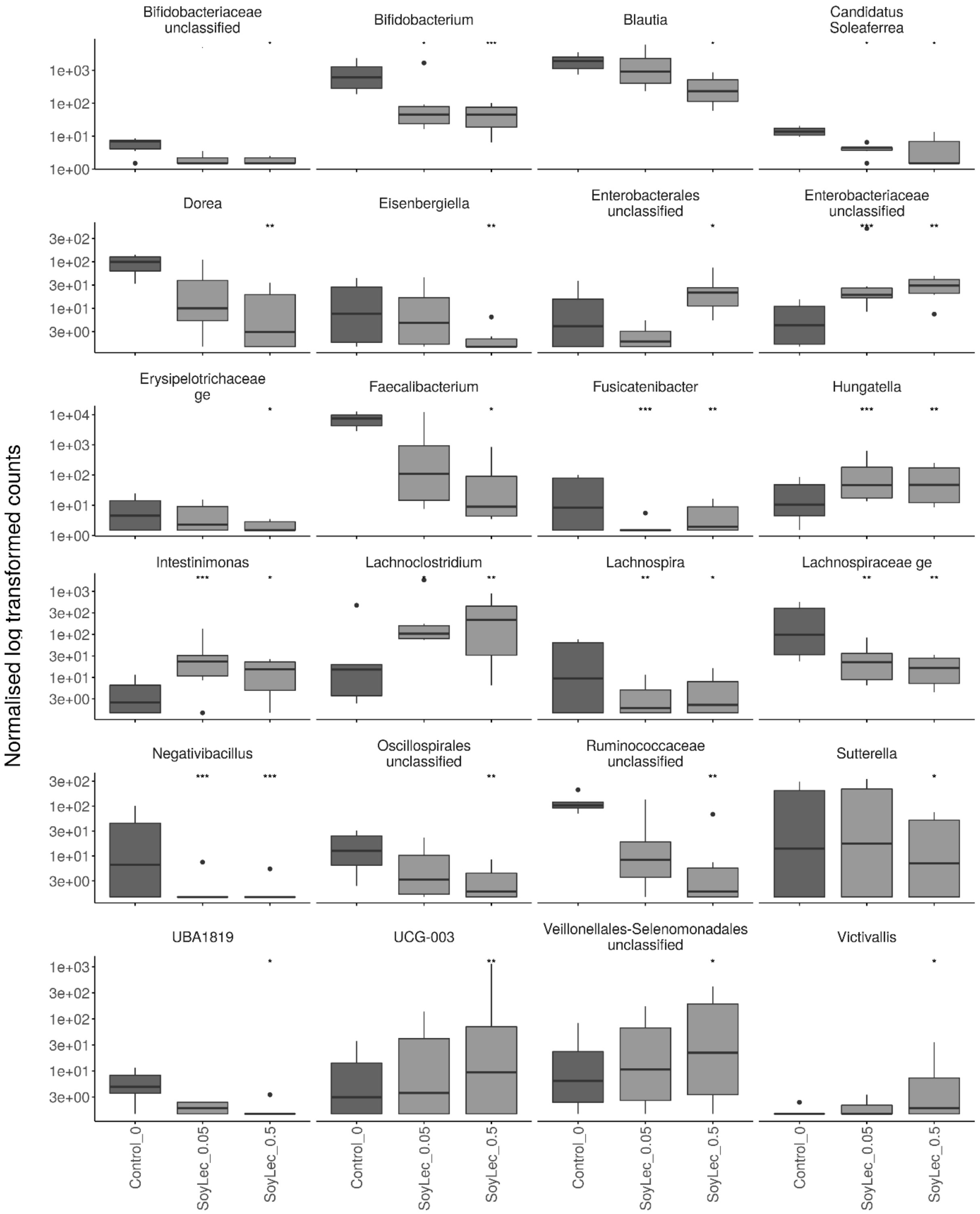


*Figure 11: Normalized counts of significantly increased or decreased genera, obtained through DESeq analysis of 16S rRNA Illumina amplicon sequencing data from luminal suspensions of a 16 and 18 day SHIME experiment investigating the impact of soy lecithin (0,05 m% and 0,5 m%) on the gut microbiota from two human faecal donors. Asterisks represent significant differences with the control based on a Wald test (α = 0,05).*

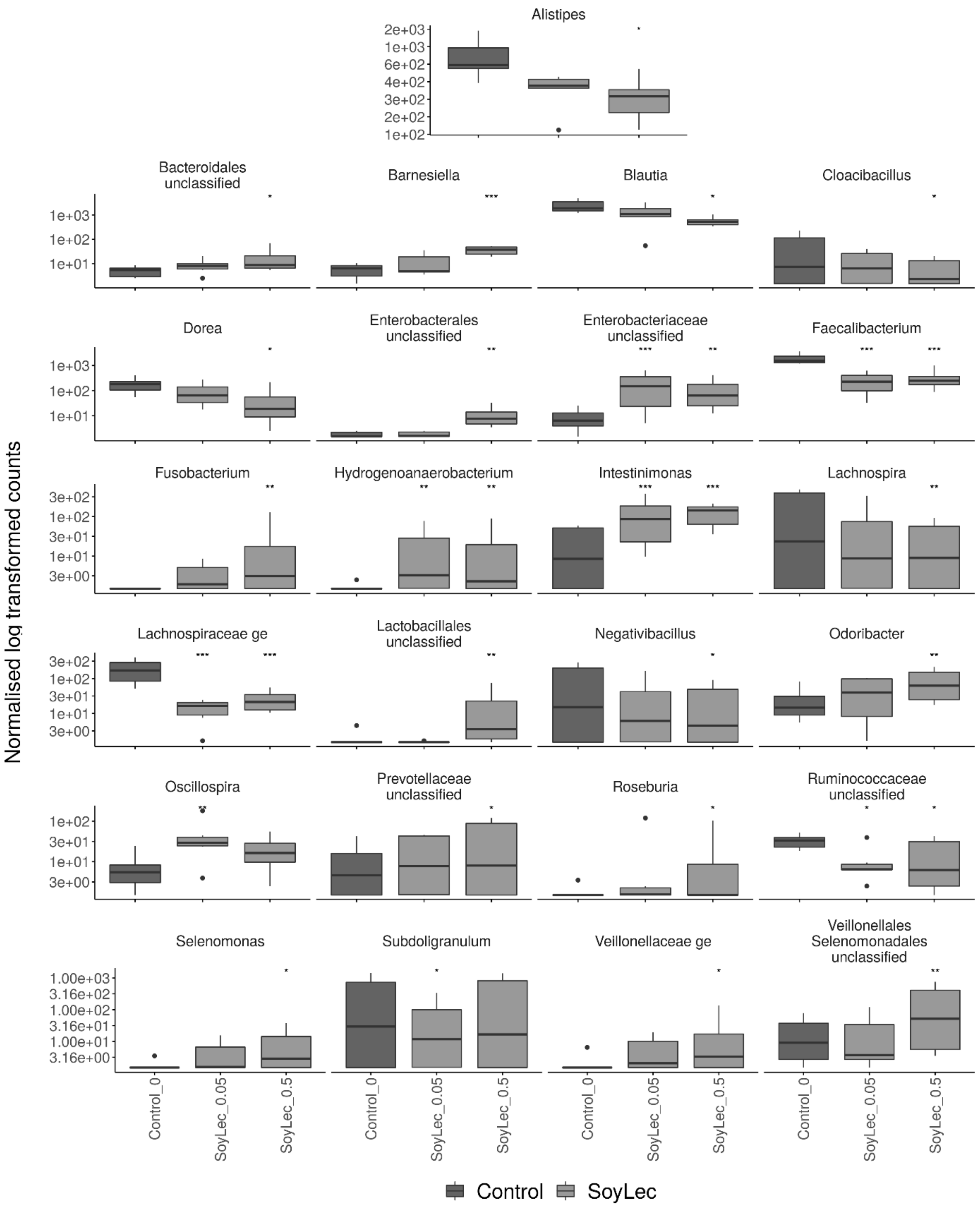


*Figure 12: Normalized counts of significantly increased or decreased genera, obtained through DESeq analysis of 16S rRNA Illumina amplicon sequencing data from mucosal compartments in a 16 and 18 day SHIME experiment investigating the impact of soy lecithin (0,05 m% and 0,5 m%) on the gut microbiota from two human faecal donors. Asterisks represent significant differences with the control based on a Wald test (α = 0,05).*

*Table 1: Log2FoldChange (L2FC) values and P-values of Wald test during DESeq-analysis of 16S rRNA Illumina amplicon sequencing data from luminal suspensions of a 16 and 18 day SHIME-experiment investigaging the impact of soy lecithin (0,05 m% and 0,5 m%) on the gut microbiota from two human faecal donors.*

| **LFC SoyLec 0.05 m%** | **LFC SoyLec 0.5 m%** | **Genus** | **P-value SoyLec 0.05 m%** | **P-value SoyLec 0.5 m%** |
|---|---|---|---|---|
| -1,76 | -3,27 | *Bifidobacterium* | 0,098 | **0,033** |
| -1,64 | -1,51 | Candidatus_Soleaferrea | 0,034 | **0,000** |
| 2,74 | 2,39 | Enterobacteriaceae_unclassified | 1,000 | **0,024** |
| -3,35 | -2,04 | *Fusicatenibacter* | **0,018** | **0,022** |
| 2,32 | 1,67 | *Hungatella* | 0,605 | **0,003** |
| 2,80 | 1,46 | *Intestinimonas* | 1,000 | **0,003** |
| 2,23 | 2,48 | *Lachnoclostridium* | 0,139 | **0,031** |
| -2,26 | -1,79 | *Lachnospira* | **0,001** | **0,003** |
| -2,43 | -2,43 | Lachnospiraceae_ge | 1,000 | **0,022** |
| -2,84 | -2,75 | *Negativibacillus* | 0,429 | **0,016** |
| -1,00 | -1,47 | Bifidobacteriaceae_unclassified | **0,000** | **0,004** |
| -0,01 | -1,71 | *Blautia* | **0,000** | **0,009** |
| -0,25 | -2,57 | *Dorea* | **0,000** | **0,033** |
| -0,03 | -2,27 | *Eisenbergiella* | 0,016 | **0,004** |
| -0,86 | 1,47 | Enterobacterales_unclassified | **0,003** | **0,013** |
| -0,05 | -1,66 | Erysipelotrichaceae_ge | 0,001 | **0,001** |
| -0,46 | -2,19 | *Faecalibacterium* | **0,000** | **0,001** |
| -0,34 | -1,96 | Oscillospirales_unclassified | 0,281 | **0,004** |
| -0,81 | -2,50 | Ruminococcaceae_unclassified | 0,156 | **0,004** |
| 0,01 | -1,33 | *Sutterella* | 1,000 | **0,032** |
| -0,86 | -1,37 | UBA1819 | 0,124 | **0,043** |
| 0,17 | 2,32 | UCG-003 | 0,679 | **0,003** |
| 0,13 | 1,70 | Veillonellales-Selenomonadales_unclassified | 0,701 | **0,014** |
| 0,02 | 1,61 | *Victivallis* | 1,000 | **0,031** |

*Table 2: Log2FoldChange (L2FC) values and P-values of Wald test during DESeq-analysis of 16S rRNA Illumina amplicon sequencing data from mucosal suspensions of a 16 and 18 day SHIME-experiment investigating the impact of soy lecithin (0,05 m% and 0,5 m%) on the gut microbiota from two human faecal donors.*

| LFC SoyLec 0.05 m% | LFC SoyLec 0.5 m% | Genus | P-value SoyLec 0.05 m% | P-value SoyLec 0.5 m% |
|---|---|---|---|---|
| 3,92 | 2,28 | Enterobacteriaceae_unclassified | **0,000** | **0,009** |
| -2,75 | -2,26 | *Faecalibacterium* | **0,000** | **0,000** |
| 2,25 | 2,04 | *Hydrogenoanaerobacterium* | **0,009** | **0,009** |
| 2,67 | 2,52 | *Intestinimonas* | **0,001** | **0,001** |
| -3,51 | -2,59 | Lachnospiraceae_*ge* | **0,000** | **0,000** |
| 2,21 | 0,80 | *Oscillospira* | **0,005** | 0,178 |
| -1,28 | -1,18 | Ruminococcaceae_unclassified | **0,049** | **0,048** |
| -1,42 | -0,02 | *Subdoligranulum* | **0,044** | 1,000 |
| -0,55 | -1,33 | *Alistipes* | 0,169 | **0,014** |
| 0,12 | 1,26 | Bacteroidales_unclassified | 0,951 | **0,049** |
| 0,18 | 2,43 | *Barnesiella* | 0,655 | **0,000** |
| -0,09 | -1,60 | *Blautia* | 0,985 | **0,014** |
| -0,61 | -1,82 | *Cloacibacillus* | 0,199 | **0,012** |
| -0,09 | -1,67 | *Dorea* | 0,985 | **0,020** |
| 0,00 | 2,26 | Enterobacterales_unclassified | 0,985 | **0,003** |
| 0,17 | 2,53 | *Fusobacterium* | 0,869 | **0,002** |
| -0,51 | -1,85 | *Lachnospira* | 0,208 | **0,005** |
| -0,05 | 2,06 | Lactobacillales_unclassified | 0,985 | **0,009** |
| -0,69 | -1,35 | *Negativibacillus* | 0,158 | **0,027** |
| 0,13 | 1,63 | *Odoribacter* | 0,688 | **0,003** |
| 0,11 | 1,16 | Prevotellaceae_unclassified | 0,891 | **0,048** |
| 0,74 | 1,45 | *Roseburia* | 0,199 | **0,048** |
| 0,28 | 1,47 | *Selenomonas* | 0,567 | **0,047** |
| **0,28** | 1,82 | Veillonellaceae_ge | 0,567 | **0,019** |
| **0,02** | 2,12 | Veillonellales-Selenomonadales_unclassified | 0,985 | **0,003** |

#### 3. Short chain fatty acids

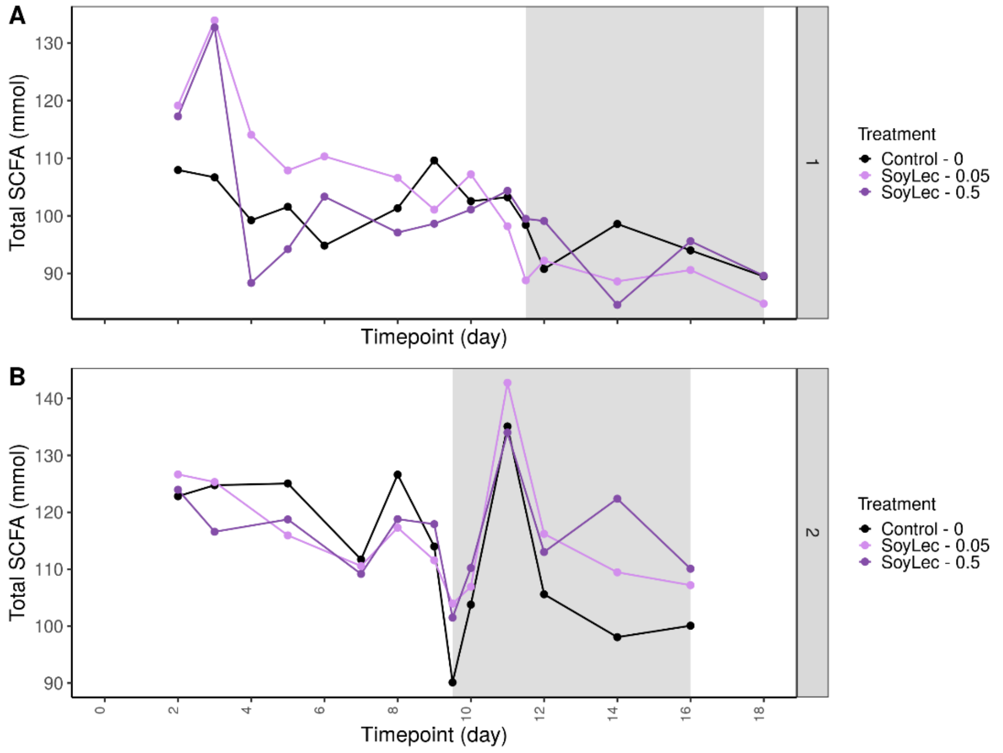


*Figure 13: Total SCFA levels (mM) in luminal suspension during a 16 and 18 day SHIME experiment investigating the impact of soy lecithin (0,05 m% and 0,5 m%) on the gut microbiota of two human faecal donors. Donor 1 was selected for low emulsifier sensitivity and donor 2 for high emulsifier sensitivity. The 7-day treatment period is indicated by the grey background.*

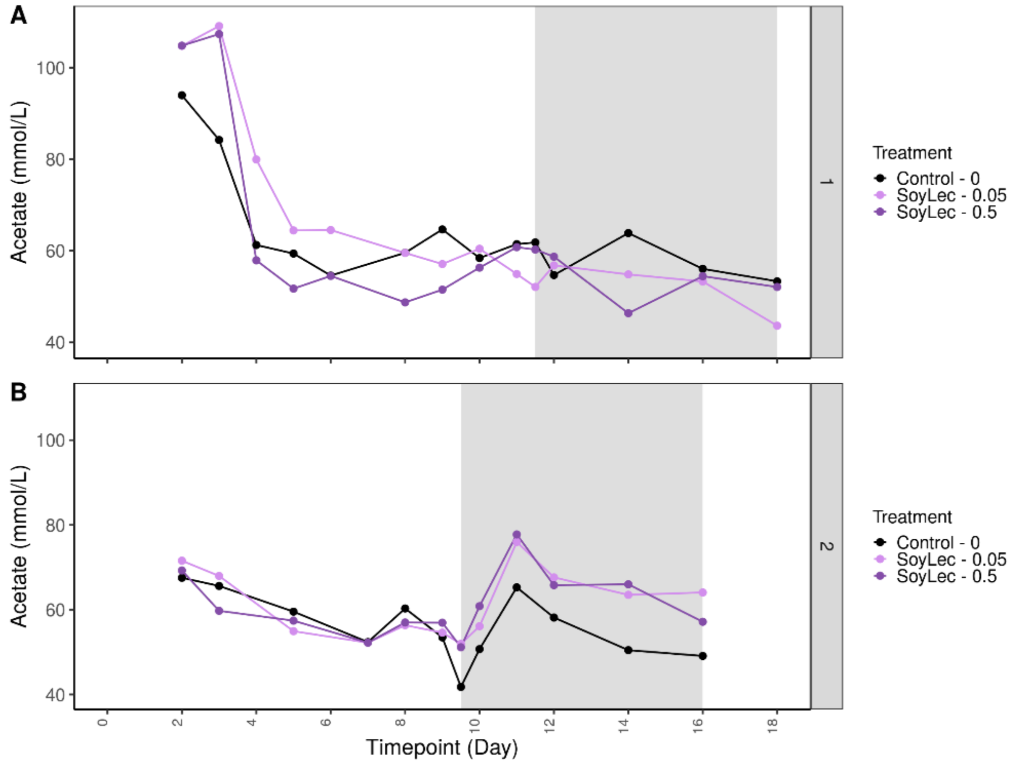


*Figure 14: Acetate levels (mM) in luminal suspension during a 16 and 18 day SHIME experiment investigating the impact of soy lecithin (0,05 m% and 0,5 m%) on the gut microbiota of two human faecal donors. Donor 1 was selected for low emulsifier sensitivity and donor 2 for high emulsifier sensitivity. The 7-day treatment period is indicated by the grey background.*

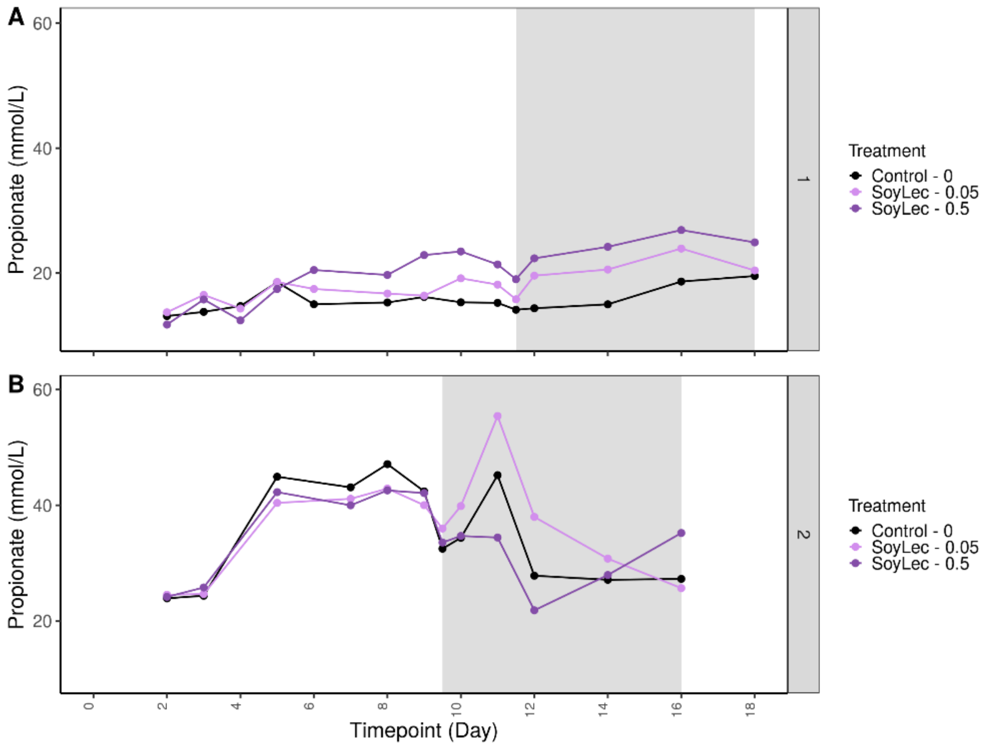


*Figure 15: Propionate levels (mM) in luminal suspension during a 16 and 18 day SHIME experiment investigating the impact of soy lecithin (0,05 m% and 0,5 m%) on the gut microbiota of two human faecal donors. Donor 1 was selected for low emulsifier sensitivity and donor 2 for high emulsifier sensitivity.The 7-day treatment period is indicated by the grey background.*

### 4. Metabolomics

#### 4.1 Targeted

##### 4.1.1 Soylec – 0,05%

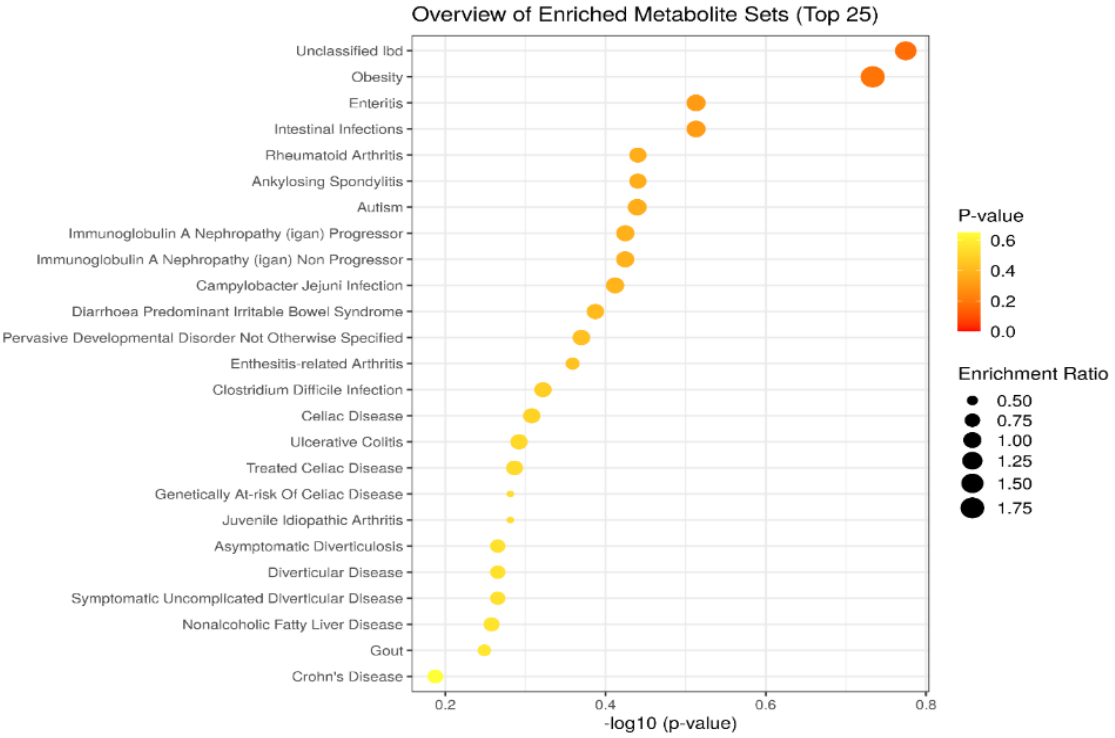


*Figure 16: Dot plot ranking disease features enriched in the human gut microbiota from two human faecal donors after exposure to 0,05 m% soy lecithin for 7 days in the Mucosal Simulator of the Human Intestinal Microbial Ecosystem (M-SHIME). Graph was obtained using the enrichment analysis tool on the MetaboAnalyst website. Disease features with a –log10 value over 1,3 are considered significant.*

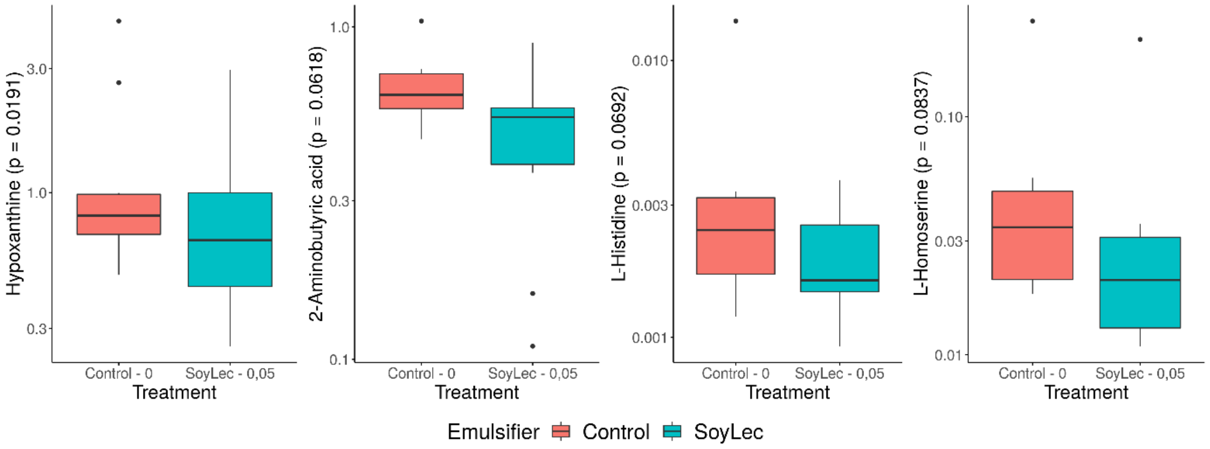


*Figure 17: Metabolites from the targeted metabolomics database annotated by the Enrichment Analysis tool on MetaboAnalyst as pointing towards IBD for the treatment with 0,05% of soy lecithin. Metabolites were detected in luminal suspension from a 16 and 18 day SHIME-experiment investigating the impact of soy lecithin (0,05 m% and 0,5 m%) on the gut microbiota of two human faecal donors during a 7 day treatment period. Samples were from timepoint 2, 11, 12, 14 and 18 from SHIME 1 and timepoint 2, 9, 10, 12 and 16 from SHIME 2.*

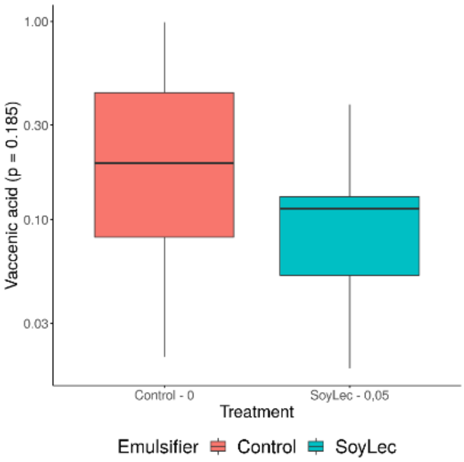


*Figure 18: Metabolites from the targeted metabolomics database annotated by the Enrichment Analysis tool on MetaboAnalyst as pointing towards obesity for the treatment with 0,05% of soy lecithin. Metabolites were detected in luminal suspension from a 16 and 18 day SHIME-experiment investigating the impact of soy lecithin (0,05 m% and 0,5 m%) on the gut microbiota of two human faecal donors during a 7 day treatment period. Samples were from timepoint 2, 11, 12, 14 and 18 from SHIME 1 and timepoint 2, 9, 10, 12 and 16 from SHIME 2.*

*Figure 19: Metabolites detected with targeted metabolomics analysis that were significantly affected by treatment of human gut microbiota with 0,05% soy lecithin in a 16 and 18 day SHIME-experiment investigating the impact of soy lecithin (0,05 m% and 0,5 m%) on the gut microbiota of two human faecal donors. Metabolites were considered significantly affected when $p_{wilcox} < 0,05$, VIP > 1 for the OPLS-DA and |p[1]| > 0,05 and |p(corr)[1]| > 0,4 on the S-plots.*

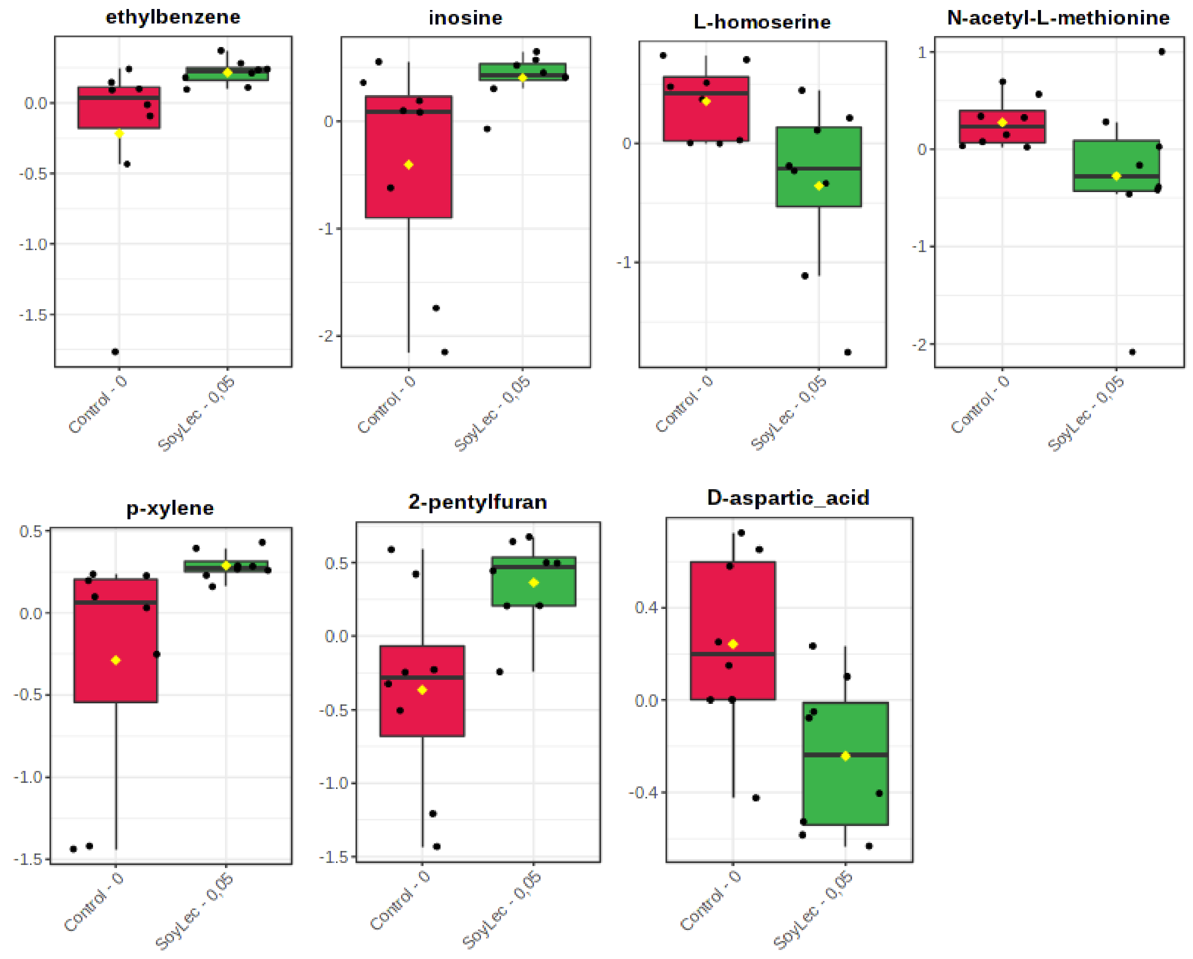


| | p_value | p[1] | p(corr)[1] | VIP |
|---|---|---|---|---|
| **p-xylene** | 0,001865 | 0,73 | 0,46 | 1,4082 |
| **ethylbenzene** | 0,014763 | 0,62 | 0,45 | 1,0492 |
| **L-homoserine** | 0,020668 | -1,16 | -0,64 | 1,7324 |
| **D-aspartic_acid** | 0,020668 | -0,67 | -0,56 | 1,181 |
| **2-pentylfuran** | 0,020668 | 1,00 | 0,56 | 1,7853 |
| **inosine** | 0,028127 | 1,02 | 0,45 | 1,9684 |
| **N-acetyl-L-methionine** | 0,049883 | -0,86 | -0,45 | 1,3421 |

##### 4.1.2 Soylec – 0,5 m%

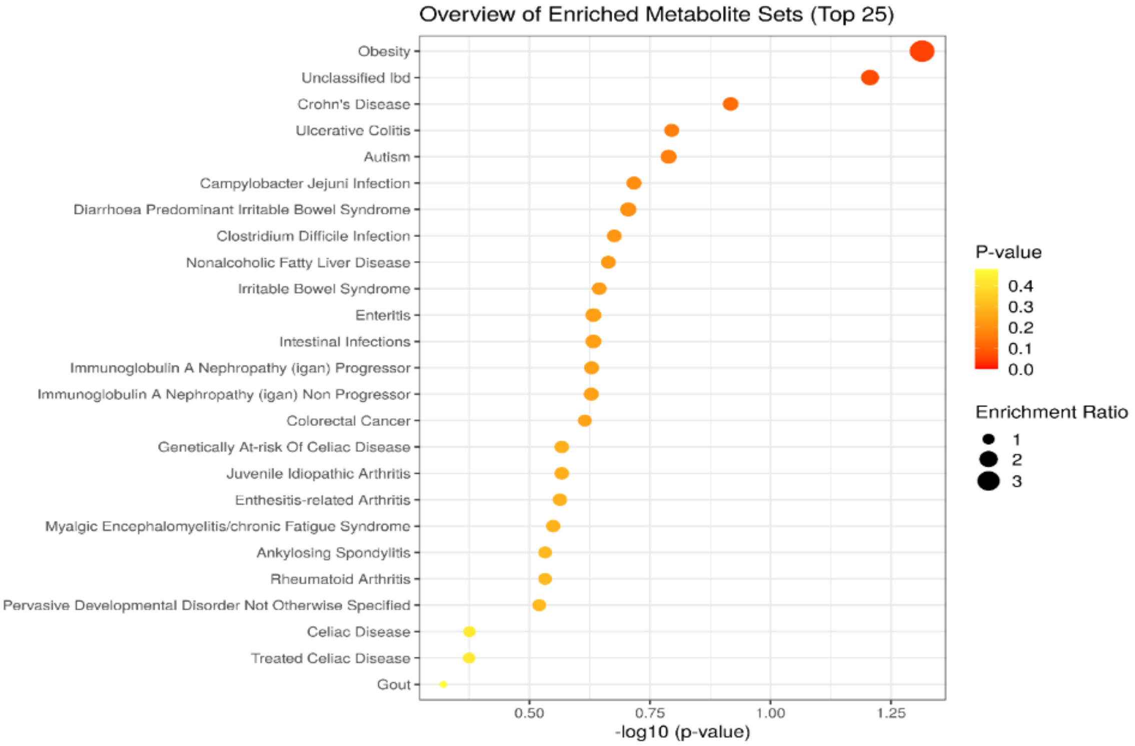


*Figure 20: Dot plot ranking disease features enriched in the human gut microbiota from two human faecal donors after exposure to and 0,5 m% soy lecithin for 7 days in the Mucosal Simulator of the Human Intestinal Microbial Ecosystem (M-SHIME). Graph was obtained using the enrichment analysis tool on the MetaboAnalyst website. Disease features with a –log10 value over 1,3 are considered significant.*

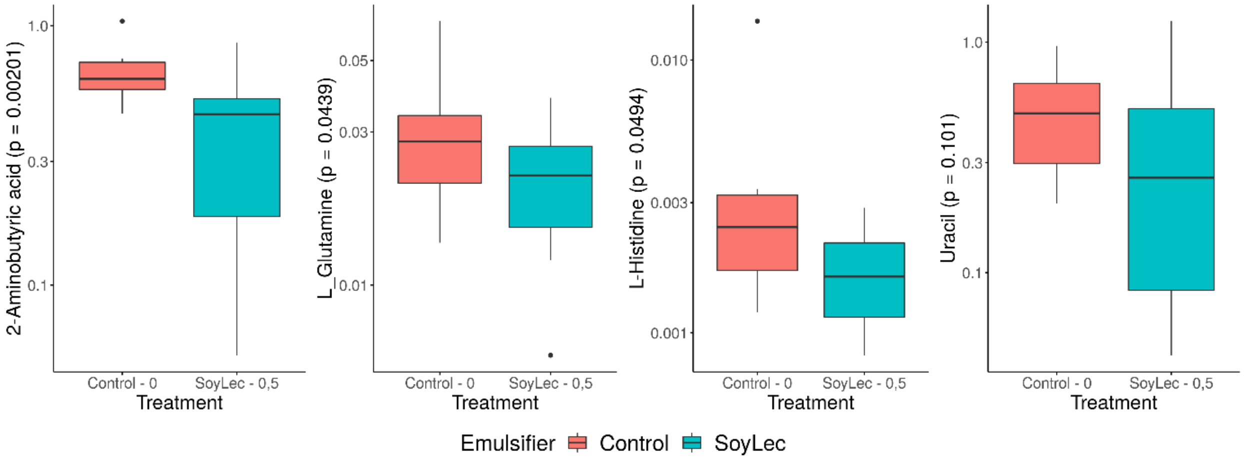


*Figure 21: Metabolites from the targeted metabolomics database annotated by the Enrichment Analysis tool on MetaboAnalyst as pointing towards IBD for the treatment with 0,5% of soy lecithin. Metabolites were detected in luminal suspension from a 16 and 18 day SHIME-experiment investigating the impact of soy lecithin (0,05 m% and 0,5 m%) on the gut microbiota of two human faecal donors during a 7 day treatment period. Samples were from timepoint 2, 11, 12, 14 and 18 from SHIME 1 and timepoint 2, 9, 10, 12 and 16 from SHIME 2.*

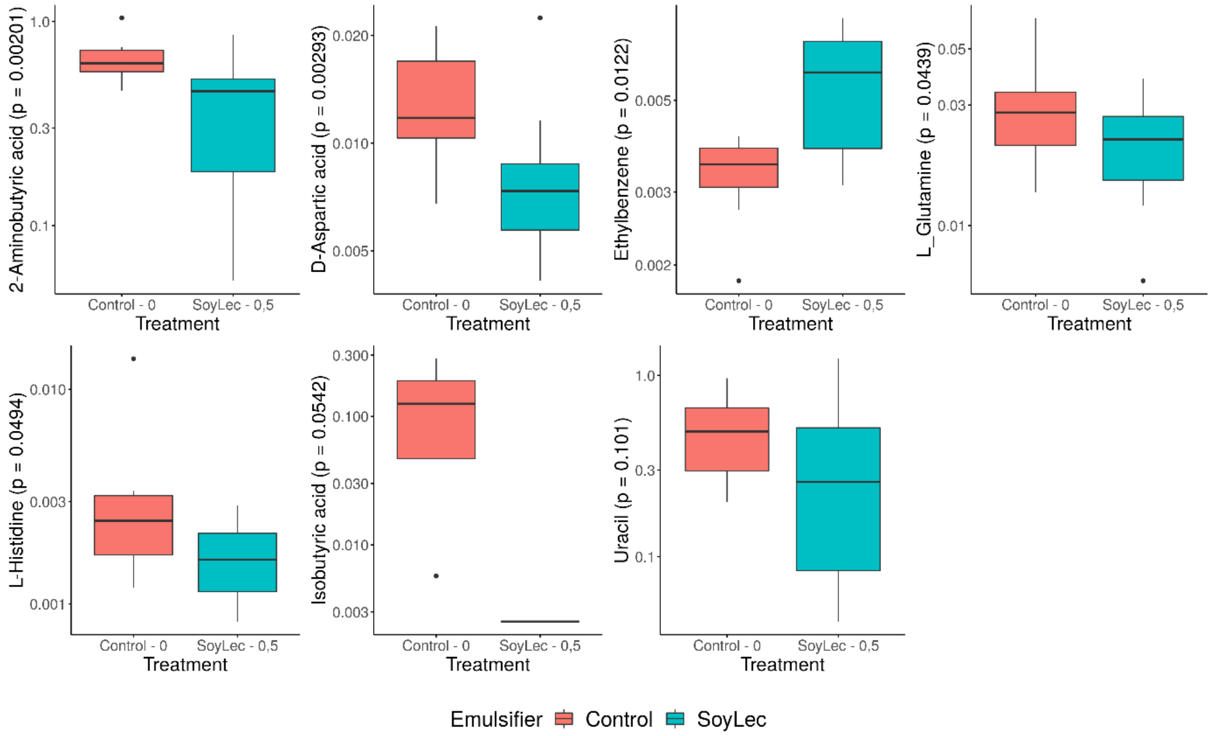


*Figure 22: Metabolites from the targeted metabolomics database annotated by the Enrichment Analysis tool on MetaboAnalyst as pointing towards crohn's disease for the treatment with 0,5% of soy lecithin. Metabolites were detected in luminal suspension from a 16 and 18 day SHIME-experiment investigating the impact of soy lecithin (0,05 m% and 0,5 m%) on the gut microbiota of two human faecal donors during a 7 day treatment period. Samples were from timepoint 2, 11, 12, 14 and 18 from SHIME 1 and timepoint 2, 9, 10, 12 and 16 from SHIME 2.*

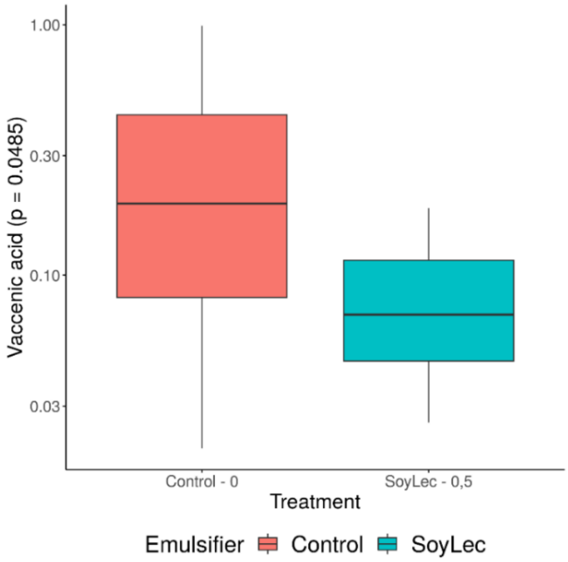


*Figure 23: Metabolites from the targeted metabolomics database annotated by the Enrichment Analysis tool on MetaboAnalyst as pointing towards obesity for the treatment with 0,5% of soy lecithin. Metabolites were detected in luminal suspension from a 16 and 18 day SHIME-experiment investigating the impact of soy lecithin (0,05 m% and 0,5 m%) on the gut microbiota of two human faecal donors during a 7 day treatment period. Samples were from timepoint 2, 11, 12, 14 and 18 from SHIME 1 and timepoint 2, 9, 10, 12 and 16 from SHIME 2.*

*Figure 24: Metabolites detected with targeted metabolomics analysis that were significantly affected by treatment of human gut microbiota with 0,5% soy lecithin in a 16 and 18 day SHIME-experiment investigating the impact of soy lecithin (0,05 m% and 0,5 m%) on the gut microbiota of two human faecal donors. Metabolites were considered significantly affected when $p_{wilcox}$ < 0,05, VIP > 1 for the OPLS-DA and |p[1]| > 0,05 and |p(corr)[1]| > 0,4 on the S-plots.*

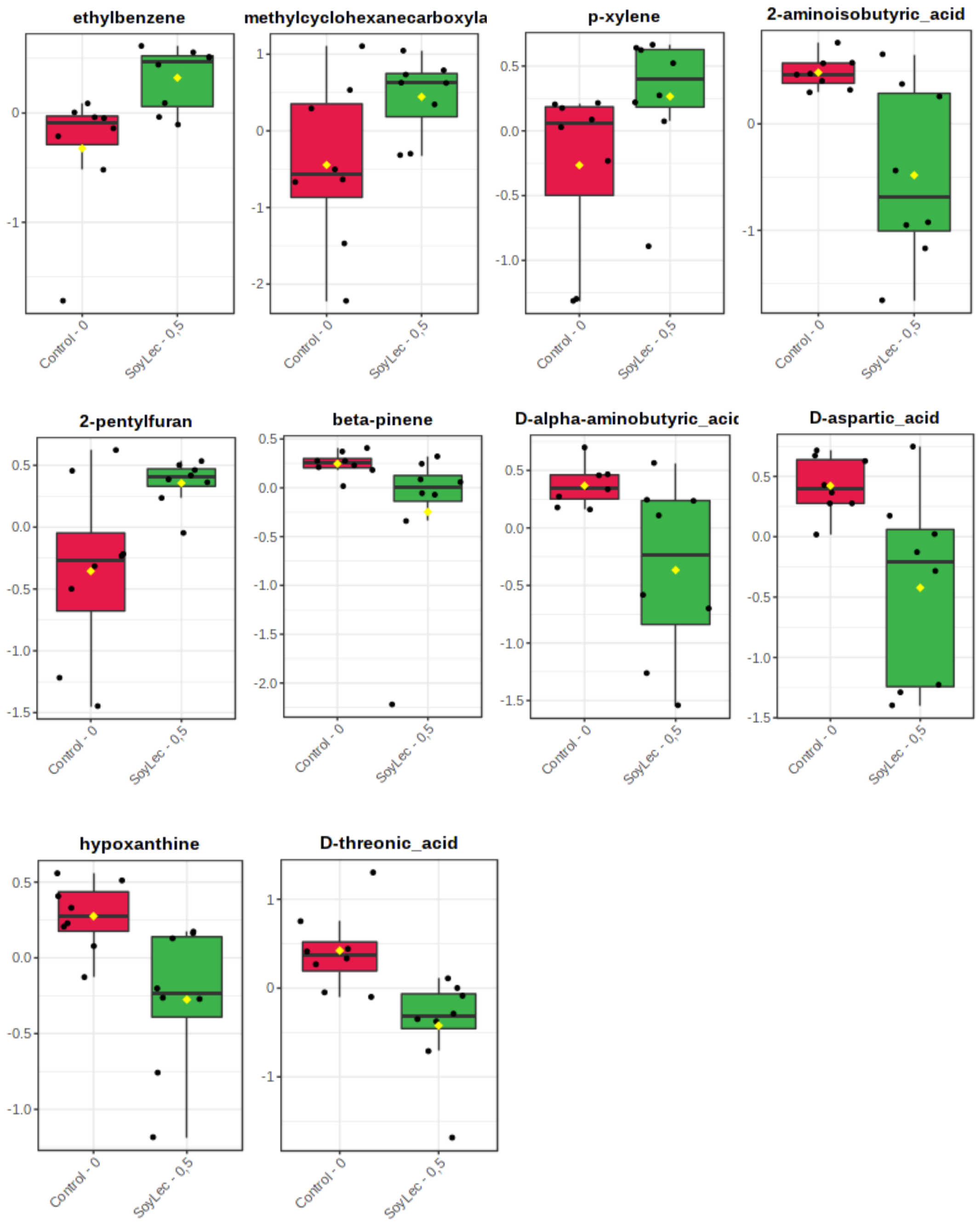


| | p.value | p[1] | p(corr)[1] | VIP |
|---|---|---|---|---|
| **D-threonic_acid** | 0,0029526 | -1,3845 | -0,65385 | 1,7627 |
| **hypoxanthine** | 0,004662 | -0,91732 | -0,6188 | 1,1497 |
| **ethylbenzene** | 0,004662 | 1,0663 | 0,59215 | 1,348 |
| **2-aminoisobutyric_acid** | 0,014763 | -1,5734 | -0,64259 | 2,0149 |
| **D-aspartic_acid** | 0,020668 | -1,4109 | -0,61642 | 1,765 |
| **p-xylene** | 0,020668 | 0,89508 | 0,44051 | 1,1086 |
| **D-alpha-aminobutyric_acid** | 0,028127 | -1,2249 | -0,5784 | 1,5299 |
| **beta-pinene** | 0,037918 | -0,85079 | -0,42585 | 1,0301 |
| **2-pentylfuran** | 0,049883 | 1,1723 | 0,58283 | 1,4874 |
| **methylcyclohexanecarboxylate** | 0,049883 | 1,5007 | 0,4997 | 1,8577 |

#### 4.2 Untargeted

*Tabel 3: Significantly different metabolites detected by Compound Discoverer (3.0) in the luminal SHIME-suspension from a 16 and an 18 days SHIME-experiment (7 day treatment period) investigating the impact of soy lecithin on the composition and functionality of the gut microbiota from two human faecal donors. Metabolites were considered significantly different when they were indicated as significantly different by the Wilcoxon Rank-sum test, the VIP value was over 1 in PLS-DA and if p(1) > 0,1 and p(corr)[1] > 0,4 in OPLS-DA for treatment with both concentrations with soy lecithin.*

| Primary ID | m/Z | RT | **Annotated name** | **Annotated Structure** |
|---|---|---|---|---|
| **4387** | 151,0688 | 2,013 | No result | No result |
| **1168** | 134,0767 | 1,352 | 3-Mercaptohexanol | C6 H14 O S |
| **2385** | 359,2591 | 14,419 | No result | C20 H38 Cl N O2 |
| **8803** | | | Myxalamid A | C26 H41 N O3 |
| **5902** | 159,931 | 3,155 | No result | No result |
| **1167** | 390,1806 | 10,999 | No result | C19 H27 N4 O3 P |

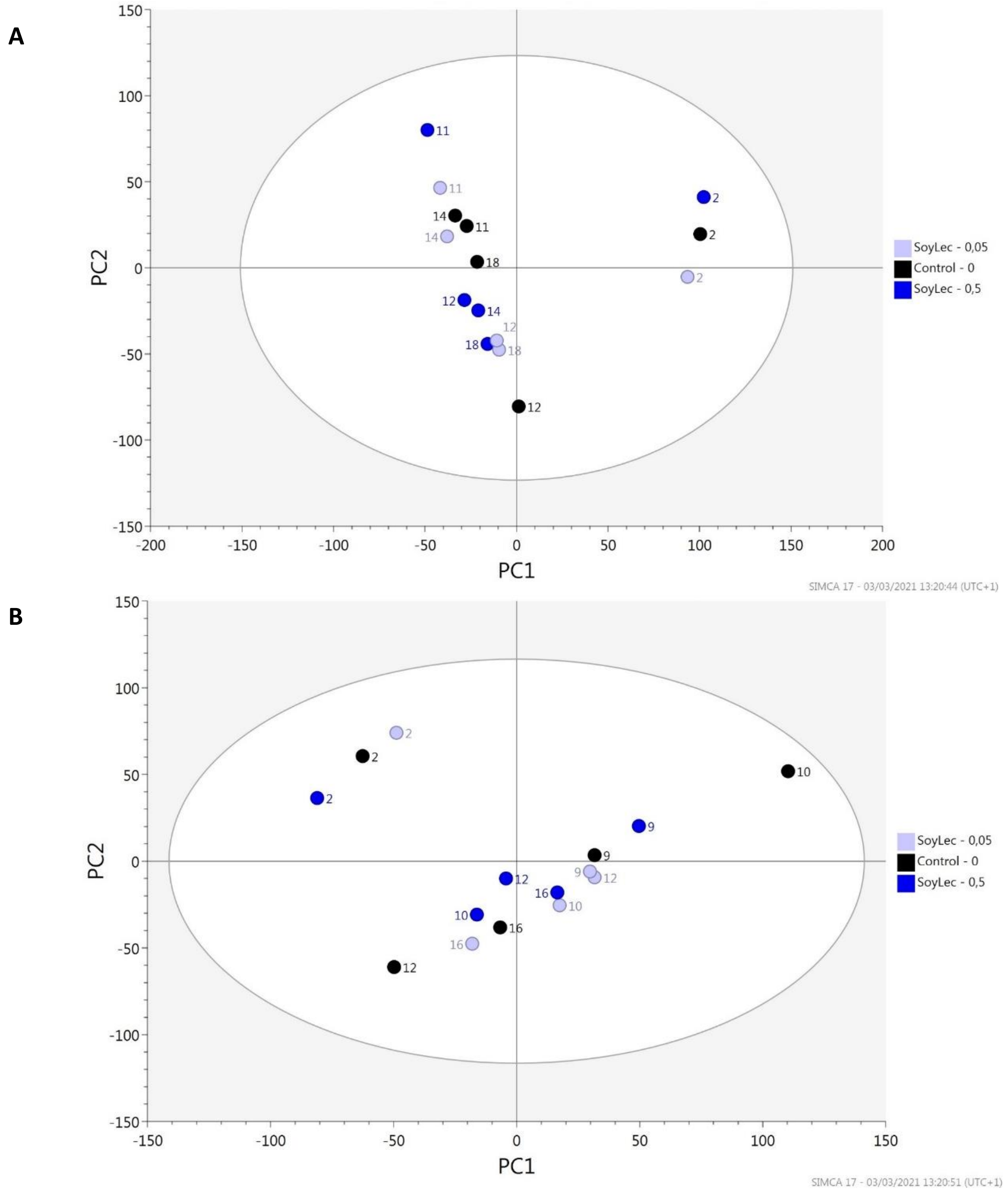


*Figure 25: Principle component analysis of untargeted metabolomics data extracted from luminal suspensions from a 16 and an 18 day M-SHIME-experiment investigating the impact of soy lecithin (0,05 m% and 0,5 m%) on the gut microbiota from two human faecal donors. A: donor 1, selected for low emulsifier sensitivity; B: donor 2, selecter for high emulsifier sensitivity.*

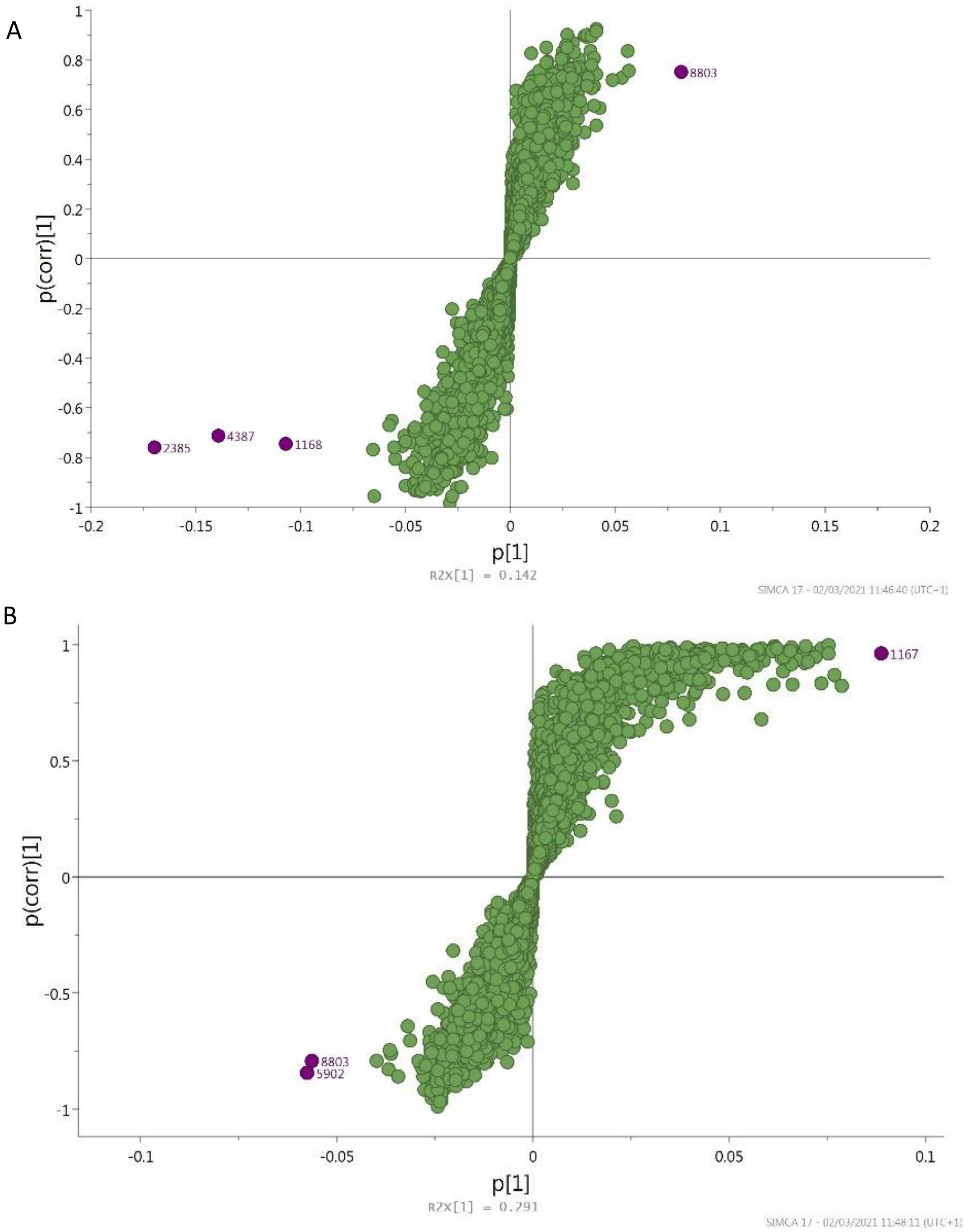


*Figure 26: S-plots obtained via orthogonal partial least squares analysis (OPLS-DA) on untargeted metabolomics data of luminal suspensions from a 16 and an 18 day M-SHIME experiment investigating the impact of soy lecithin (0,05 m% and 0,5 m%) on the composition and functionality of the gut microbiota from two human faecal donors. The OPLS-DA models compared both soy lecithin treatments – 0,05 m%: A and 0,05 m% : B - with the control. P[1] represents the relative magnitude of the change. P(corr)[1] represents the confidence/reliability of the effect. As a result the data points in the upper right and lower left corners are the least likely the result of spurious correlations (T. Zhang et al., 2015). All data points designate certain metabolites. Metabolites were considered potential biomarkers (purple) when p[1] > 0,05 and p(corr)[1] > 0,4.*
